# Quantitative single-cell analysis of *Leishmania major* amastigote differentiation demonstrates variably extended expression of the lipophosphoglycan (LPG) virulence factor in different host cell types

**DOI:** 10.1101/2022.06.27.497826

**Authors:** Michael A. Mandell, Wandy L. Beatty, Stephen M. Beverley

**Author notes:** Corresponding author; Dept. of Molecular Microbiology, Washington University School of Medicine, Campus Box 8230, 660 S. Euclid Ave., St. Louis MO 63110.

## Abstract

Immediately following their deposition into the mammalian host by an infected sand fly vector, *Leishmania* parasites encounter and are engulfed by a variety of cell types. From there, parasites may transit to other cell types, primarily macrophages or dendritic cells, where they replicate and induce pathology. During this time, *Leishmania* cells undergo a dramatic transformation from the motile non-replicating metacyclic stage to the non-motile replicative amastigote stage, a differentiative process that can be termed amastigogenesis. To follow this at the single cell level, we identified a suite of experimental ‘landmarks’ delineating different stages of amastigogenesis qualitatively or quantitatively, including new uses of amastigote-specific markers that showed interesting cellular localizations at the anterior or posterior ends. We compared amastigogenesis in synchronous infections of peritoneal and bone-marrow derived macrophages (PEM, BMM) or dendritic cells (BMDC). Overall, the marker suite expression showed an orderly transition post-infection with similar kinetics between host cell types, with the emergence of several amastigote traits within 12 hours, followed by parasite replication after 24 hours, with parasites in BMM or BMDC initiating DNA replication more slowly. Lipophosphoglycan (LPG) is a *Leishmania* virulence factor that facilitates metacyclic establishment in host cells but declines in amastigotes. Whereas LPG expression was lost by parasites within PEM by 48 hours, >40% of the parasites infecting BMM or BMDC retained metacyclic-level LPG expression at 72 hr. Thus *L. major* may prolong LPG expression in different intracellular environments, thereby extending its efficacy in promoting infectivity *in situ* and during cell-to-cell transfer of parasites expressing this key virulence factor.

**Author Summary:** Leishmaniasis caused by species of the trypanosomatid protozoan parasite *Leishmania* is an important widespread global health problem. Humans become infected by *Leishmania* following the bite of sand flies bearing the flagellated metacyclic promastigote stage, and the first 24-48 hours thereafter are critical to the outcome of infection. During this time, parasites are engulfed by several host cell types, where they differentiate into the rounded amastigote stage, adapted for intracellular survival and proliferation, with concomitant changes in metabolism and virulence factor expression. We developed a suite of markers that allowed us to monitor promastigote-to-amastigote differentiation (amastigogenesis) on a single-cell level. Two showed amastigote-specific expression and localized to the anterior or posterior regions. Our marker suite allowed us to chart the course of amastigogenesis in different host cell environments and to determine the timing of amastigote development relative to the initiation of parasite replication and the expression of the virulence factor lipophosphoglycan (LPG). We report that the amastigogenesis process follows a determined sequence of events that occurs prior to parasite replication. In contrast, parasites may respond to different host cell environments by prolonging LPG expression, thereby extending the duration of its pro-infectivity functions within or in transit between host cell destinations.

## Introduction

The establishment phase of infection following the initial encounter with the host is a crucial time for parasites, as it is during this time that an often-small number of colonizing organisms are either successfully transmitted allowing for replication or are eliminated. Pathogens transmitted by arthropod vectors face additional challenges in that they must transition from an insect host-adapted to a mammalian host-adapted stage of the life cycle. *Leishmania* parasites are transmitted to their mammalian hosts by the bite of an infected sand fly which deposits non-replicating, flagellated metacyclic-stage parasites into wounded dermis while taking a blood meal [1]. In order to establish a successful infection, parasites must overcome many obstacles. First, they must survive lysis by complement [2] and entanglement in neutrophil nets [3, 4] while in the extracellular milieu. Next, the parasites must encounter and be engulfed by a phagocytic cell. Once inside, a parasite must convert the host cell into an environment suitable for replication [5], differentiate into the non-motile amastigote stage of the life cycle in a process that we refer to as amastigogenesis, and re-enter the cell cycle [1].

Because macrophages are the cell type most commonly infected in established *Leishmania* infections, *in vitro* studies investigating the various parasite and host processes implicated in virulence have been primarily carried out in these cells. While these studies typically employ relatively homogenous populations of macrophages, reality is more heterogeneous as the incoming population of infecting parasites individually encounter a wide variety of different cell types *in vivo*, whose full complexity of interactions is not fully understood. Most studies *in vitro* or *in vivo* emphasize the role of professional phagocytes encountering and engulfing infective metacyclic stage parasites, including various macrophage subsets, monocytes, neutrophils and dendritic cells (DCs) [6–8]. Many of these are rapidly recruited to the infection site, and collectively may predominate over macrophages as *Leishmania* host cells [6–8]. Non-immune cell types such as keratinocytes have also been observed harboring parasites *in vivo* where they may play significant roles [9, 10]. Moreover, parasites can ‘transit’ from one cell type to another, for example from neutrophils to DCs or macrophages [6–8, 11]. These cell-to-cell transitions likely happen within 1-2 days after infection, since the majority of parasites are found within macrophages after this period [7, 8, 12, 13].

Different cell types encountered by invading *Leishmania* metacyclics likely present different challenges, reflecting variation in the intracellular environment and innate immune responses. How the parasites differentiate within and respond to varying cell-type specific environments is not completely understood. In order to overcome the obstacles inherent in becoming established, *Leishmania* parasites employ an array of temporally regulated virulence factors which have important roles early in infection by metacyclic-stage parasites but are down-regulated later in infection [14, 15]. One such virulence factor is the promastigote surface glycoconjugate, lipophosphoglycan (LPG) [5]. LPG is the most abundant molecule on the surface of invading metacyclic-stage parasites [16], and protects *Leishmania* parasites from killing by complement and neutrophil extracellular traps [4, 17]. LPG also has important roles in rendering infected cells safe niches for parasite survival and replication [18]. LPG is shed into the infected host cell where it interferes with phagosome biogenesis [19], oxidant defenses [17], signal transduction pathways [20], antigen presentation [21], and transiently inhibits phagolysosomal fusion and acidification [22, 23]. Despite these important roles for LPG in early infection, its synthesis is reduced in amastigote-stage parasites, possibly to avoid activation of Caspase-11 or other immune regulators [24].

Here, we tested whether the nature of the host cell encountered by *L. major* metacyclic stage parasites has consequences for the parasite in terms of its ability to differentiate into the amastigote stage and commence replication. To this end, we identified temporally spaced experimental “landmarks” that clearly distinguish metacyclics from amastigotes and allow microscopic visualization of the process of amastigogenesis at the single cell level. Interestingly, two new marker epitopes were found which additionally exhibited striking cellular localization at the anterior and posterior amastigote ends, which may warrant further study in the future. Using these “landmarks” in synchronous infections, we found that parasite differentiation proceeded during the first 12 hours after infection in all cell types tested with an orderly sequence of marker transitions. Despite synchronous expression of amastigote markers, parasite re-entry into the cell cycle differed significantly between host cell types. Parasites within cells that supported reduced parasite replication prolonged expression of LPG relative to parasites within the more highly permissive cell types. Together, these data show that different host cell types present different intracellular environments and that *Leishmania* can respond to these different environments by modulating the duration of virulence factor expression.

## Results

### Characterization of differentiation markers and their expression following metacyclic L. major infection

We first sought markers that clearly distinguished *L. major* metacyclic parasites from authentic amastigotes, and which could serve as useful temporal ‘landmarks’ to assess amastigogenesis following infection across populations of cells microscopically following synchronous infections. While multiple genes showing quantitative differences in promastigote or amastigote expression are known from microarray or proteomic studies [25–31], few show qualitatively on/off properties suitable for microscopic evaluation of the dynamics of amastigote differentiation on a cellular level. Candidate markers were evaluated using comparisons of purified infecting metacyclic-stage parasites versus replicating amastigotes (72 hours after infection of PEMs; Fig. 1). In these experiments, metacyclics were allowed to attach/invade for only 2 hours prior to removal of unbound parasites, in order to have a relatively synchronous infection. We identified six useful markers, four of which show qualitatively on/off properties. As a marker that is ‘on’ in procyclic and metacyclic promastigotes and transitions to ‘off’ in amastigotes, we chose the expression of the paraflagellar rod protein PFR2, which is lost along with the flagellum during differentiation (Fig. 1, top row) [32, 33].

**Figure 1.**
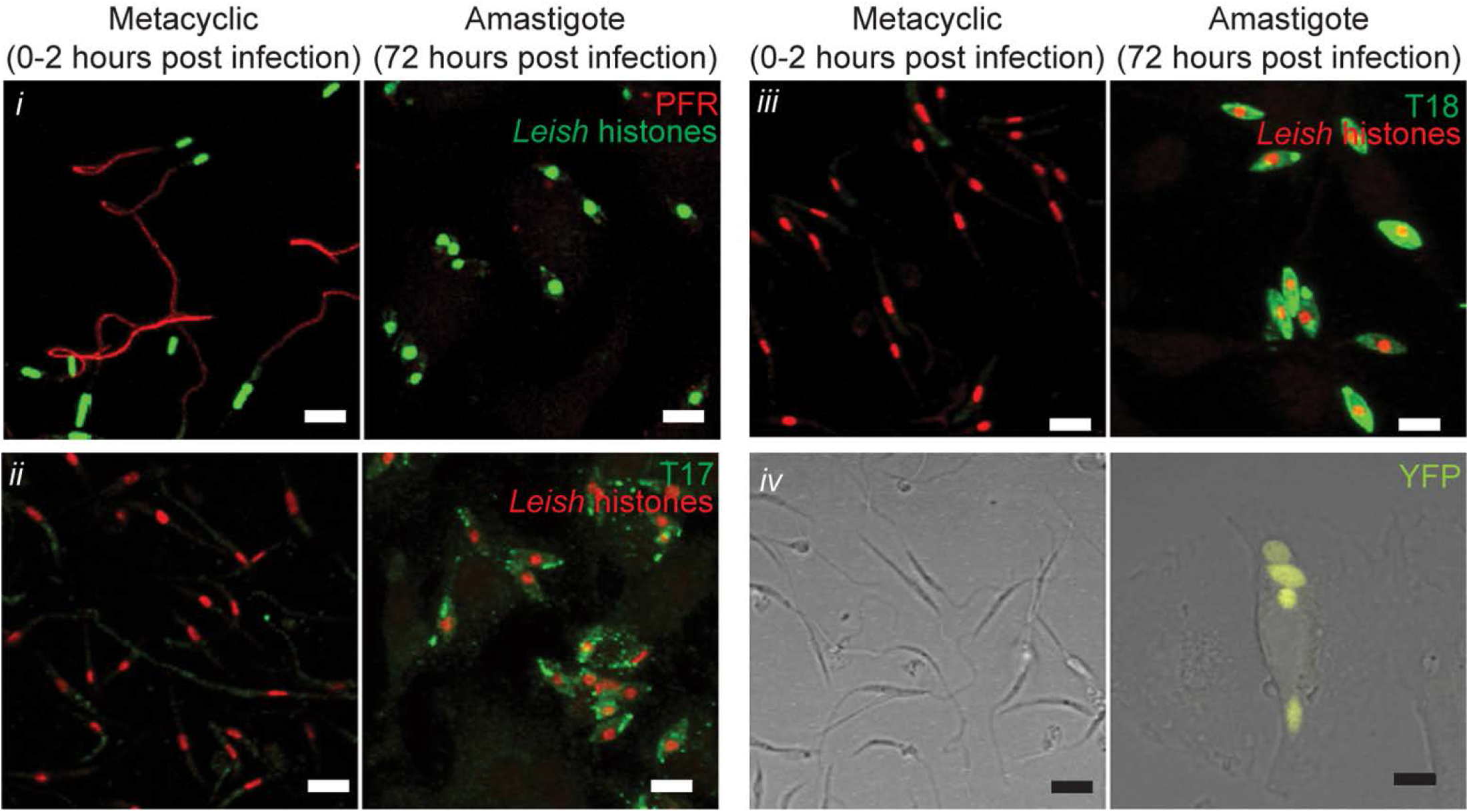
Cytological markers differentiate metacyclic promastigote from amastigote stage *Leishmania major.* Comparison of marker expression on metacyclic promastigotes (left) or parasites 72 hours post infection of PEMs (i.e. amastigotes, right). (*i*) Parasite nuclei are detected with antisera against *L. major* histone proteins (green), and PFR is shown in red. (*ii*) Parasite histones, red. mAB T17, green. (*iii*) Parasite histones, red. mAB T18, green. (*iv*) YFP fluorescence (yellow) overlaid onto DIC image. Scale bar represents 5 µm.

As markers ‘off’ in promastigotes and ‘on’ in amastigotes, we selected expression of the amastigote-specific antigens recognized by the monoclonal antibodies T17 and T18, which emerged from studies seeking stage or species-specific tools [34]. Both antisera yielded a strong labelling of amastigotes but not incoming metacyclics (Fig. 1, rows 2 and 3). The molecular identity of the epitopes recognized by T17 and T18 are unknown, but amastigote localization data suggest they represent independent molecular entities as discussed in a later section.

A fourth marker consisted of a YFP transgene inserted into the ribosomal SSU locus, where YFP is “off” in metacyclic promastigotes but “on” in other stages (Fig. 1, bottom row). YFP fluorescence is lost as parasites enter stationary growth phase (Fig. S1A); it is during this stage of *in vitro* growth that a subset of parasites differentiate into the metacylic stage [35, 36]. Stationary-phase parasites express less YFP at both the protein and mRNA level (Fig. S1B, C) demonstrating that YFP is down-regulated at the transcriptional level, although the precise mechanism has not been studied. As discussed below, YFP fluorescence is restored in most, but not all, amastigotes.

### Morphological changes associated with amastigogenesis

We additionally established two quantitative markers for amastigogenesis: loss of the metacyclic flagellum and the transition to the more rounded amastigote morphology. We first assessed the disappearance of the parasite flagella (Fig. 2A, B). Because PFR2 expression is lost very rapidly following infection, other markers were used to detect flagella. At early time points, we used an antibody against the parasite glycocalyx component lipophosphoglycan (LPG), which allowed visualization of the entire parasite surface (Fig. 2A, left panel). However, as LPG expression is eventually lost following infection of mammalian cells, at later time points over 8 hours we visualize parasite flagella using the T17 marker (Fig. 2A, right panel). We then determined what fraction of parasites had “long” flagella (LF; ≥ 5 µm; Fig. 2B). As expected, 100% of metacyclic stage parasites had LF prior to infection. Two hours after infection, the overwhelming majority (89 ± 3%) of parasites retained LF. However, by 8 hours after infection the fraction of parasites with LF had sharply declined to 21 ± 4%, and by 24 h only 6 ± 1% had LF. Similar to what was described for *L. mexicana* [37], our data indicate that the acquisition of the amastigote flagellum resulted from flagellar shortening rather than a digital switch between long and short flagella, as we measured flagellar lengths intermediate between metacyclic (11.8 ± 0.5 µm) and amastigote (2.8 ± 0.1 µm) at 8 hours post infection (4.8 ± 0.4 µm; Fig. 2C).

**Figure 2.**
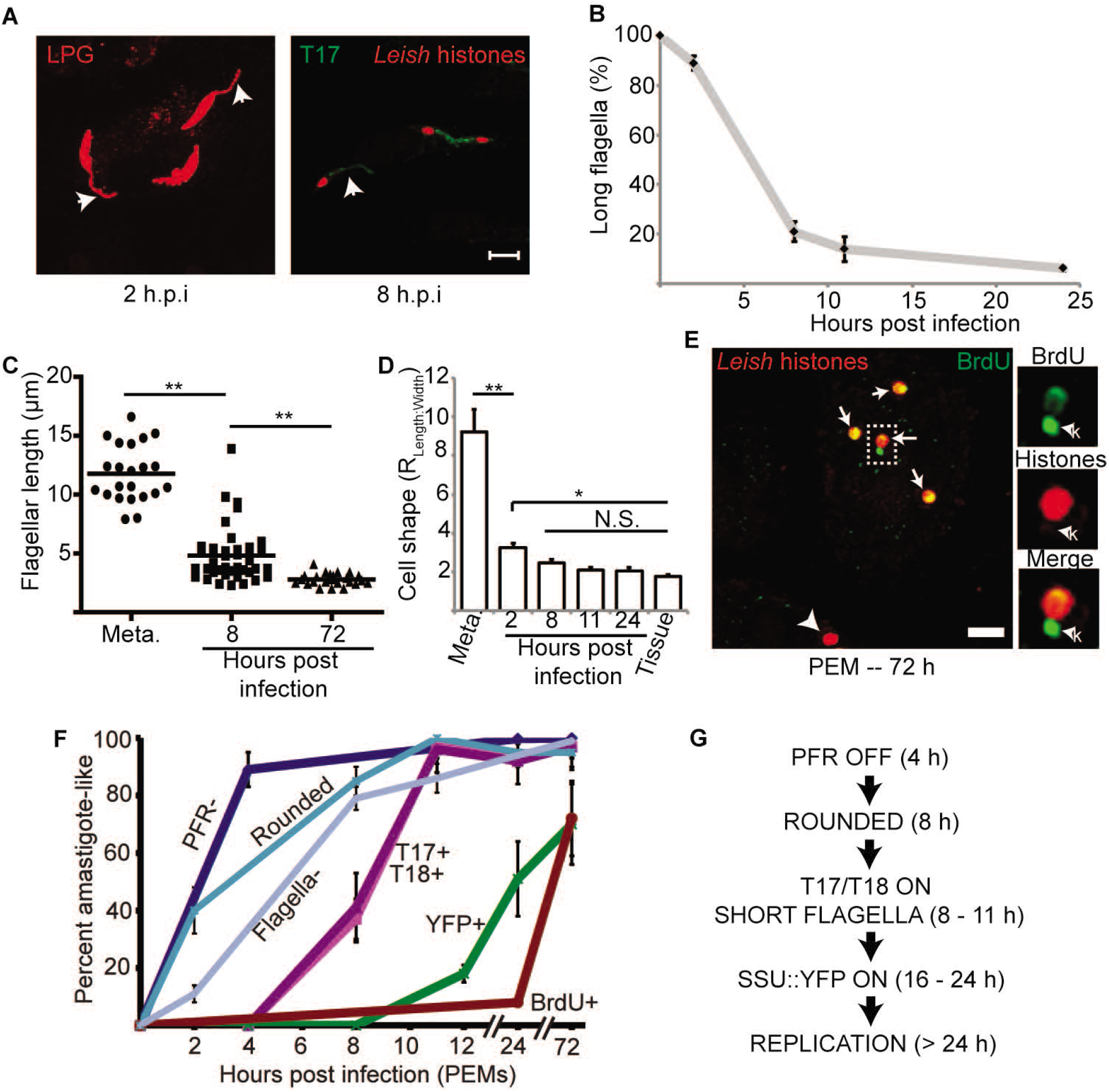
Timing of developmental changes associated with amastigote differentiation in PEMs. **(A)** Parasite flagella were identified at different time points following infection of PEMs with anti-LPG (≤ 8 h post infection) or mAb T17 at longer time points. The fraction of parasites retaining long flagella (≥ 5 µm; arrows) was determined at the indicated time points and plotted in **(B)**. Scale bar represents 5 µm. N > 100 parasites analyzed per time point. **(C)** Quantitative analysis of parasite flagellar length of metacyclics or parasites at the indicated time points post infection of PEMs. Each data point represents one parasite. **(D)** The kinetics of *L. major* metacyclic-stage parasites acquiring amastigote-like shape characteristics. Metacyclic-stage parasites, parasites at the indicated time points post infection of PEMs, or amastigotes from infected mouse footpad tissue 2 weeks post infection were stained to detect the parasite cell body and imaged by confocal microscopy. Image analysis was then performed to determine the length and width of each parasite, with cell shape determined as the ratio between parasite length and width. Metacyclic parasites and parasites within PEMs were stained with anti-LPG (time points ≤ 8 h or mAb T18 (time points > 8 h). Tissue amastigotes were detected by YFP fluorescence. N > 50 parasites. **(E)** Confocal micrograph of BrdU-labeled parasites. PEMs were infected with metacyclic-stage *L. major* for 72 h in the presence of 0.1 mM BrdU prior to fixation and immunolabeling to detect BrdU (green) and *Leishmania* histones (red). Arrows indicate parasite nuclei showing BrdU-labeling. Arrowhead, BrdU-negative parasite nucleus. Boxed region is magnified in the images on the right. k, BrdU-positive kinetoplast. Scale bar, 5 µm. **(F)** The percentage of parasites showing an “amastigote-like” phenotype for the various markers is plotted as a function of time after infection of PEMs. N > 200 parasites analyzed per marker per time point. **(G)** Schematic summary of the data in (F) showing the timing of amastigote development culminating in new DNA synthesis. Data, means ± S.E.M for 3 independent experiments; *, *P* < 0.05; **, P < 0.001 by ANOVA.

To assess parasite rounding during amastigogenesis, we determined the ratio between the length and width of the parasite cell body. For these experiments, metacyclics and parasites up to 8 h after infection were detected with anti-LPG antibodies, while parasites at later time points were visualized with T18 antibody. We found that *L. major* assumes an “amastigote” shape very rapidly (Fig. 2D), with parasites 2 h after infection being substantially more round than metacyclics (metacyclic length to width ratio 9.2 ± 1.1 vs 3.2 ± 0.2 two hours post infection; P < 0.001). By 8 h after infection, the average parasite shape was indistinguishable from that seen in tissue sections from mice that had been infected for 2-3 weeks.

### Sequence and timing of metacyclic/amastigote differentiation marker expression

We then performed a time course analysis of the expression of our suite of parasite differentiation markers following infection of PEMs by metacyclics (Fig. 2F). In addition to the markers described above, we determined the timing at which non-replicating metacyclics resume replication after infection by assessing BrdU incorporation as a measure of parasite DNA synthesis (Fig. 2E) [38]. The first event was a rapid and nearly quantitative loss in reactivity with anti-PFR2 antisera which preceded the disappearance of the flagella. By 4 hours post-infection (the earliest time point attempted), most parasites had lost anti-PFR staining (89 ± 6%), and by 24 hours essentially no parasites (0.2 ± 0.6%) were recognized with this antibody. These next were followed by parasite rounding and flagellar loss as described above.

The emergence of reactivity with mAbs T17 and T18 is first observed around 8 hr after infection. At four hours post-infection, only a small percentage of parasites (<5%) showed reactivity with either antibody. By 8 hours post-infection, 37 ± 7% and 41 ± 12% were strongly positive for T17 or T18 reactivity, and by 11 hours after infection, 97 ± 4% or 96 ± 8% were positive for T17 and T18 reactivity, respectively. Because both primary antibodies are raised in mice, using indirect IFA we could not simultaneously determine both markers, however the quantitative data suggest they emerge together.

The SSU:YFP transgene expression first became weakly detectable 8 hours after infection, with 18 ± 3% of the parasites YFP^+^ at 12 hours post-infection. By 24 hours, the percentage of YFP^+^ parasites had increased to 51 ± 13%. While the percent of parasites displaying YFP fluorescence continued to increase reaching 70 ± 17% at 72 hours post-infection, thereafter it never exceeded that value *in vitro* (Fig. 2F) or *in vivo* as monitored in infected mouse footpads (Fig. S2A). The reason this marker never attains 100% expression in amastigotes is unknown, but the effect seems to be transitory since 100% of promastigotes show YFP fluorescence after isolation from mouse tissue (data not shown). In dual-labeling experiments, we found that induction of T17 and T18 reactivity always preceded YFP expression at a time point at which neither marker has reached its maximum (10 hrs after infection), since at this time point 100% of YFP-positive parasites were T17 or T18 positive (Fig. S2B, C)

As judged by BrdU incorporation, DNA replication did not commence significantly until after 24 hr, well after other markers had transitioned to the amastigote state. At 24 hr, PFR expression was undetectable while T17 and T18 expression were maximal; in contrast, only 8 ± 1% of the parasites showed labeling with anti-BrdU antisera at this time, increasing to 70% at 72 hr (Fig. 2F).

Together, these data show an ordered sequence, with promastigote-specific PFR2 gene expression turning off by 4 hr post-infection, T17 and T18 expression turning on by 8-11 hr, SSU:YFP expression turning on by 11-24 hr, and finally replication commencing after 24 hr. These data establish a useful developmental sequence of marker expression for analysis of *Leishmania* amastigogenesis (Fig. 2G).

### Congruent induction profiles of amastigotigenesis markers in PEM, BMM, and BMDC

We then used the amastigogenesis marker suite to compare the timing of *L. major* development following synchronous infection of two different macrophage types (PEM, BMM) as well as DC (BMDC) *in vitro*. These cell types have been used in numerous *Leishmania* infection studies *in vitro*, and they were chosen because they allowed us to assess the diversity of parasite responses to entry into different host cell environments. Loss of anti-PFR2 reactivity in BMM and DCs was similar to what is seen in PEMs at either 4 or 24 hours post-infection with 35 ± 25% and 0.5 ± 0.6% of parasites in BMMs and 15 ± 13% and 0.4 ± 4% of parasites in DCs labeling at these time points (Fig. 3A). The induction of T18 reactivity and SSU:YFP fluorescence also showed similar time courses in the three cell types (Fig. 3B, C). Induction of T17 reactivity appeared to be slower in BMMs than in PEMs in that a lower percentage of the parasites at 8 hours post-infection were T17^+^ (10 ± 8%, P < 0.01). This difference disappeared by 24 hours, with 89 ± 10% of the parasites within BMMs showing T17-positivity. Parasites within DCs were intermediate between the profile seen for PEMs and BMMs in terms of T17 labeling at 8 hours post infection, but were not significantly different from the PEM results (Fig. 3D). We also found that parasites within BMM and BMDC were comparable to parasites in PEM in terms of cell shape 24 hours after infection (Fig. 3E).

**Figure 3.**
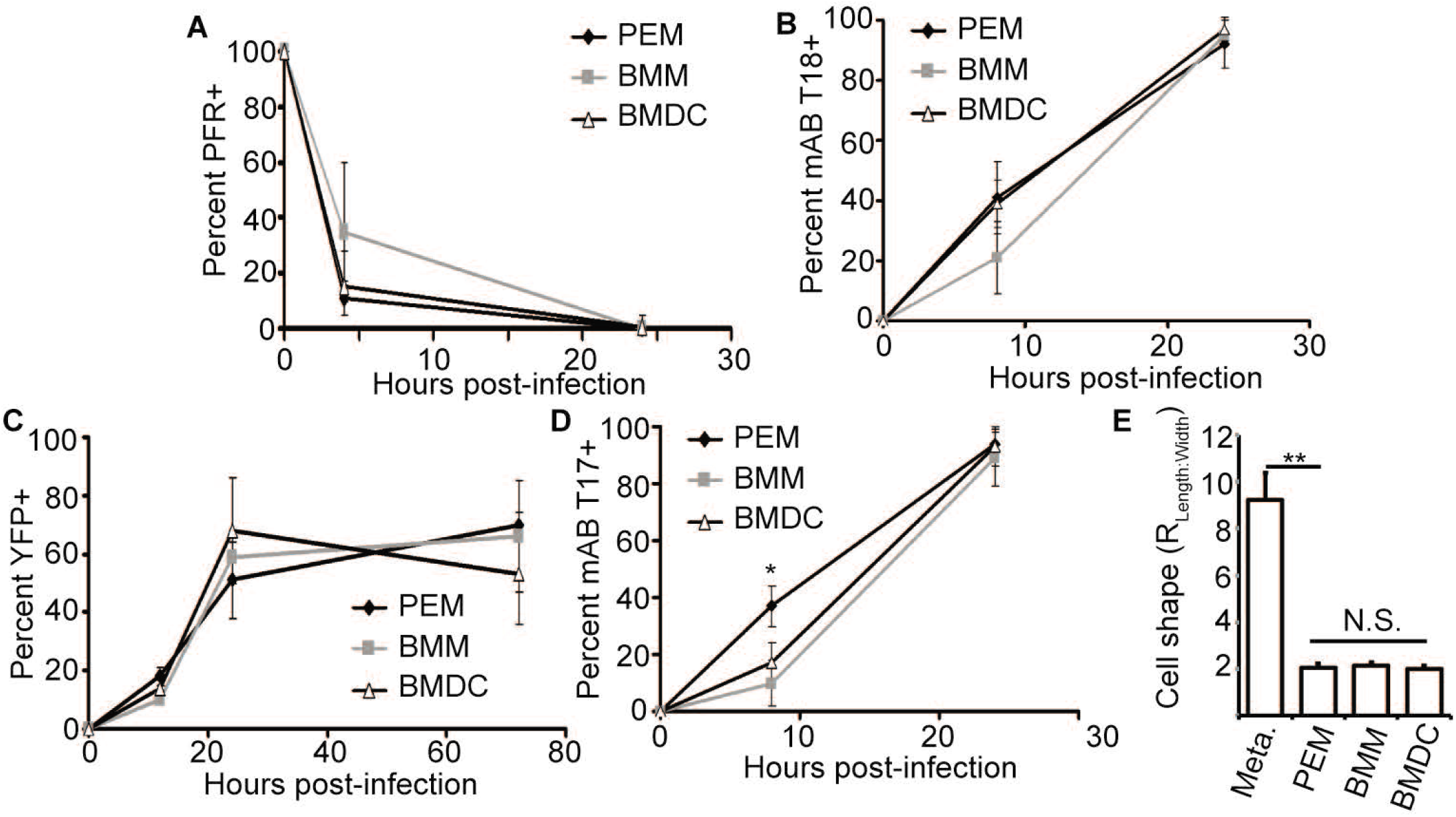
Metacyclic-to-amastigote transition is similar in different host cell types. **(A-D)** Comparison of metacyclic-to-amastigote transition in PEM (black diamonds), BMM (gray squares) and BMDC (open triangles) for the percent of parasites positive for PFR labeling (A), mAb T18 labeling (B), YFP signal (C), and mAb T17 labeling (D). **(E)** Comparison of parasite cell shape 24 h after infection of the indicated cell types with metacyclic-stage *L. major.* Data, means ± S.E.M for 3 independent experiments; *, *P* < 0.05; **, P < 0.001 by ANOVA; N.S., not significant. N > 350 parasites per marker/time point.

These data suggested the possibility of a conserved ‘core’ amastigogenesis program. To test this, we transferred stationary phase *L. major* (the time at which metacyclogenesis is maximal) from 25° C to 37° C in the absence of host cells for 24 hr. As in other *Leishmania* systems, this was sufficient to trigger amastigote marker transitions and amastigote morphology. Under these conditions, 90% of parasites showed PFR loss, 60% showed upregulation of T17 and T18 antigens, and 27% showed YFP positivity, (Fig. S3A-C). That these developmental changes also occurred in axenic culture demonstrates that early steps of amastigogenesis are largely independent of host cell type.

### Parasite replication is delayed in BMM and BMDC

We asked whether the timing of parasite re-entry into the cell cycle differed following infection of the different cell types using the BrdU incorporation assay. In all host cell types, very few parasites (5-8%) labeled with BrdU during the first 24 h post-infection (Fig. 4A). By 72 hr after infection, 35 ± 8% of parasites in BMM and 46 ± 5% of parasites in BMDC were BrdU positive, significantly less than what is seen in PEM (72 ± 7%). In agreement with these results, we observed substantial parasite replication as measured by the number of parasites per host cell in PEM but not in BM-derived cells over a 72 hr infection time course (Fig. 4B). Taken together, these data indicate that PEMs are a relatively more permissive host cell type for replication of *L. major* following metacyclic infection than BMM or BMDC.

**Figure 4.**
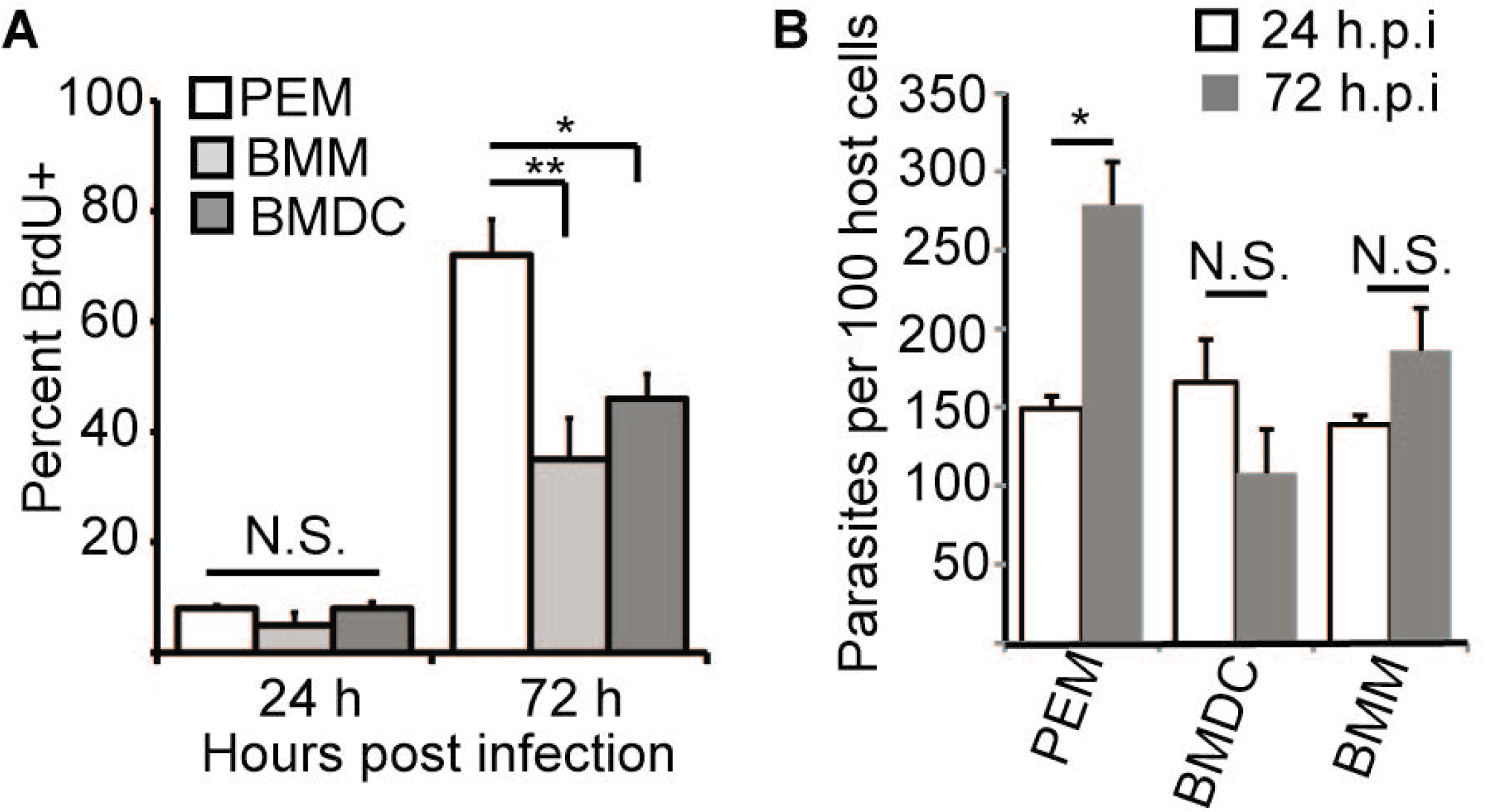
Initiation of parasite replication delayed in bone-marrow derived macrophages and DC. **(A)** PEM, BMM, and BMDC were infected with metacyclic-stage parasites and cultured in the presence of BrdU for the indicated time points prior to fixation. The percent of parasites showing BrdU-positive nuclei was determined by analysis of samples stained with anti-BrdU and anti-*L. major* histone antibodies. N > 350 parasites per cell type/time point. **(B)** The number of parasites per 100 host cells at 24 and 72 h post infection of PEM, BMM, and BMDC. Data, mean ± S.E.M. of three independent experiments. *, *P* < 0.05, **, *P* < 0.001 by ANOVA. N.S., not significant.

### Quantitation of LPG expression

Given the similarities in overall amastigogenesis in the three cell types, we then explored the timing of expression of the *Leishmania* virulence factor LPG, which normally shuts off in amastigotes. In PEM, LPG expression was assessed at different time points after infection by its reactivity with the phosphoglycan (PG) specific monoclonal antibody WIC79.3, which recognizes galactose modifications of LPG and proteophosphoglycan (PPG) [18, 39]. While these modifications may be further capped by arabinose in metacyclic LPG in *L. major*, previous studies showed metacyclic parasites purified by gradient centrifugation retain high levels of exposed Gal-residues and WIC79.3 reactivity [36]. While PPGs also react with WIC79.3, they are much less abundant than LPG (<1%) [39].

First, LPG expression was assessed qualitatively, scoring parasites as either “LPG-positive” or “LPG-negative”. By this assay, 99 ± 1% of metacyclic stage parasites were LPG^+^. By 24 hours after infection of PEMs, this number had declined to 81 ± 2%, and by 72 hours post infection, only a few (5 ± 3%) of the parasites had detectable LPG. Experiments monitoring the presence of arabinose-capped metacyclic-specific LPG using the monoclonal antibody 3F12 yielded comparable results with a modest decrease in the percentage of 3F12-positive parasites by 24 hours after infection and almost complete loss of 3F12^+^ parasites by 72 hours post infection (Fig. S4A).

The intensity of WIC79.3 reactivity was then used to quantitate LPG expression per *L. major* cell (Fig. 5A). All metacyclic parasites showed high levels of LPG, with a mean labeling intensity of 46,000 ± 2,600 arbitrary units. At two hours post infection the mean per-cell WIC79.3-reactivity increased to 66,100 ± 3,530 units (P < 0.0001), and thereafter, LPG expression gradually declined steadily over the next 24 hr. By 48 hours post infection, LPG levels had declined to background levels, as established by comparisons to the Δ*lpg1*^-^ mutant lacking LPG [18] (Fig. 5A). LPG loss occurs during the period where the amastigote marker YFP is induced, and so we tested whether these two events occurred in an ordered sequence (e.g. YFP “on” before LPG “off” or vice versa). However, quantitative analysis indicated that the timing of LPG loss appears to be random relative to YFP induction (Fig. S4B-D), suggesting that these two events do not happen in an ordered sequence.

**Figure 5.**
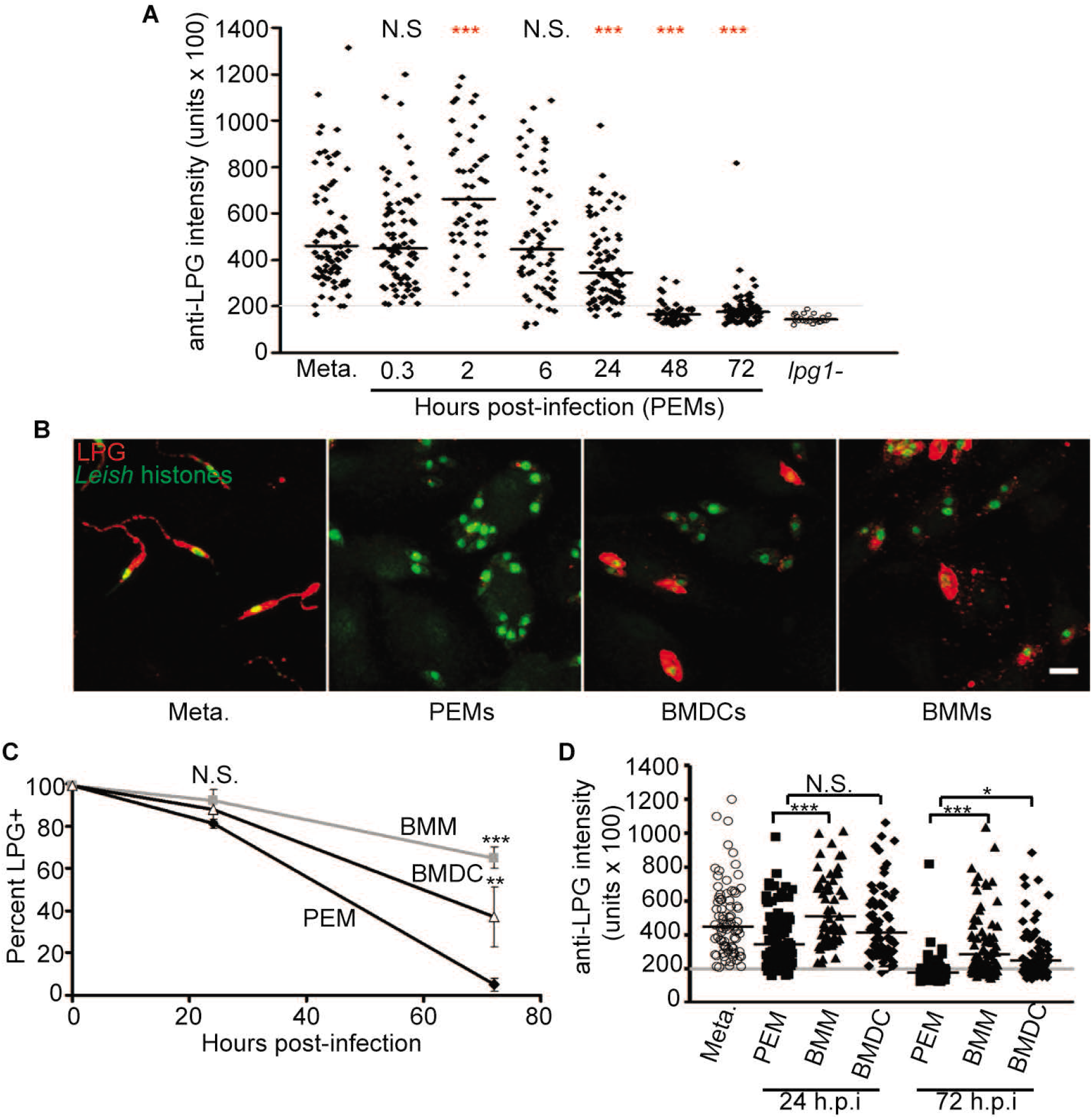
Retention of LPG expression following infection of BMMs and BMDCs. **(A)** Anti-LPG fluorescence intensity on a per-parasite basis. Anti-LPG intensity of WT metacyclic-stage parasites as well as at various time points after infection of PEMs was measured as described in Methods. As a negative control, the anti-LPG intensity of *L. major lpg1-* (open circles) was measured 0.3 hr after infection of PEMs. The gray line shows the mean anti-LPG intensity of ‘LPG-negative’ WT parasites plus two standard deviations, and any parasite with anti-LPG values below that line would be considered “LPG-negative”. Black bars represent geometric mean of the data for each sample. Data shown is pooled from at least two independent experiments. ***, *P* < 0.0001 compared to the values for metacyclics using ANOVA with Bonferroni post hoc analysis. **(B)** Representative images comparing the LPG-positivity of purified metacyclics and parasites within PEMs, DCs, and BMMs at 72 hours post-infection. Parasite nuclei are shown in green, LPG is shown in red. Scale bar represents 5 µm. **(C)** Percent of parasites within the three host cell types that are LPG^+^ as determined by the ‘qualitative’ assay. PEM data, black diamonds, BMM data, gray squares, DC data, open triangles. N > 200 parasites per time point/cell type. **(D)** Quantitation of LPG on the surface of parasites within PEMs, BMMs, and DCs at 24 and 72 hours post infection. For comparison, anti-LPG intensity data for metacyclic-stage parasites is also shown. The gray line shows the cut-off for LPG-positivity. Black bars represent geometric mean of the data for each sample. n ≥ 3 experiments. N.S, not significant, *, *P* < 0.05; **, *P* < 0.001; ***, *P* < 0.0001 (ANOVA).

### High levels of LPG are retained on some parasites for an extended period in BMMs and BMDCs

We examined LPG expression following metacyclic infection of BMM and BMDCs (Figs. 5B-D). The profiles were very similar to what was seen in PEMs for the first 24 hr, with few parasites showing LPG-negativity in the three host cell types (Fig. 5C). However, by 72 hours after infection, a striking difference emerged. Whereas 95 ± 3% of parasites within PEMs were qualitatively LPG-negative at 72 hours after infection, in both BMMs and BMDCs a much higher percentage of parasites retained LPG expression with 65 ± 5% of parasites in BMMs (*P* < 0.0001) and 37 ± 14% of parasites in BMDCs retaining LPG (*P* < 0.001) (Fig. 5B, C). Accordingly, fluorescence intensity measurement showed that the LPG-positive subset of parasites at late time points expressed quantitatively robust levels of LPG within BMM and BMDC (Fig. 5D). At 24 hours after infection, the mean LPG intensity of parasites within PEMs had declined by ∼25% with 7% of the cells showing staining intensity comparable to background. In contrast, LPG expression on parasites within BMM and BMDC was not different from what was seen on metacyclics with essentially no LPG-negative parasites detected. By 72 hours post infection, 78% of parasites in PEM had background levels of LPG staining and the mean anti-LPG intensity had gone down by ∼60% (17,600 ± 900 units versus 44,800 ± 2,300 units for metacyclics). While the LPG staining intensity of parasites within BMM and BMDC did decline by the 72 hour time point, these parasites retained significantly more LPG (28,600 ± 2,100 and 24,900 ± 1,700 mean anti-LPG intensity, respectively) than did parasites in PEM. Notably, we observed a great deal of heterogeneity between *L. major* cells within the BMM and BMDC. While some parasites (∼35%) were LPG-negative and resembled those seen in PEMs, ∼15% of the parasites retained levels of LPG expression as high as that seen in metacyclic-stage parasites (>43,400 units). Together, these data show that *L. major* can alter the duration of LPG expression in response to different host cell environments with a population of parasites retaining biologically relevant amounts of LPG for at least 72 hours after infection.

### LPG-retaining parasites within bone-marrow dendritic cells and macrophages have not initiated DNA synthesis

We asked whether the strongly LPG^+^ sub-populations of infected BMM or BMDC cells after 72 hr had initiated DNA synthesis. We used the Click-iT EdU system (Life Technologies) [40] to facilitate simultaneous visualization of LPG and the incorporation of thymidine analogue (EdU) into parasite DNA. BMDCs were infected with metacyclics and then cultured in the presence of EdU for 72 hours, after which the samples were fixed and stained to simultaneously detect *L. major* histones, LPG, and EdU. Importantly, the results obtained using EdU were similar to the results described above with BrdU. We imaged a total of 1174 parasites, 44% of which were EdU^+^ and 21% of which were LPG^+^ (Fig. 6A, B). While 54% of LPG-negative parasites were EdU^+^, only 7% of the LPG-positive parasites were EdU^+^, significantly less than what would be expected if LPG- and EdU-positivity behaved independently of each other (*P* < 10^-30^ by a Chi-square analysis; Fig. 6B). Similar results were obtained when BMM were tested following BrdU labeling (Fig. S5A, B). This demonstrates that *L. major* parasites in BMDC and BMM have a strong tendency to down-regulate LPG expression prior to cell cycle re-entry. Next, we asked if LPG loss occurred in an ordered fashion and always preceded initiation of DNA synthesis. To assess this, we performed the same analysis as above on parasites within PEM at 24 hours post infection, a time point at which most parasites retain LPG expression yet a few parasites have become BrdU-positive. Unlike in BMM and BMDC at 72 hours, LPG-positivity of EdU-positive parasites was essentially random with LPG/EdU double positive parasites readily observed (Fig. S5C, D). A summary comparing the course of events following parasite entry of PEM, BMDC, and BMM is shown in Figure 6C. Overall, our data indicate that LPG loss is not a prerequisite for parasite replication, but there is a strong tendency for non-replicating parasites to retain LPG.

**Figure 6.**
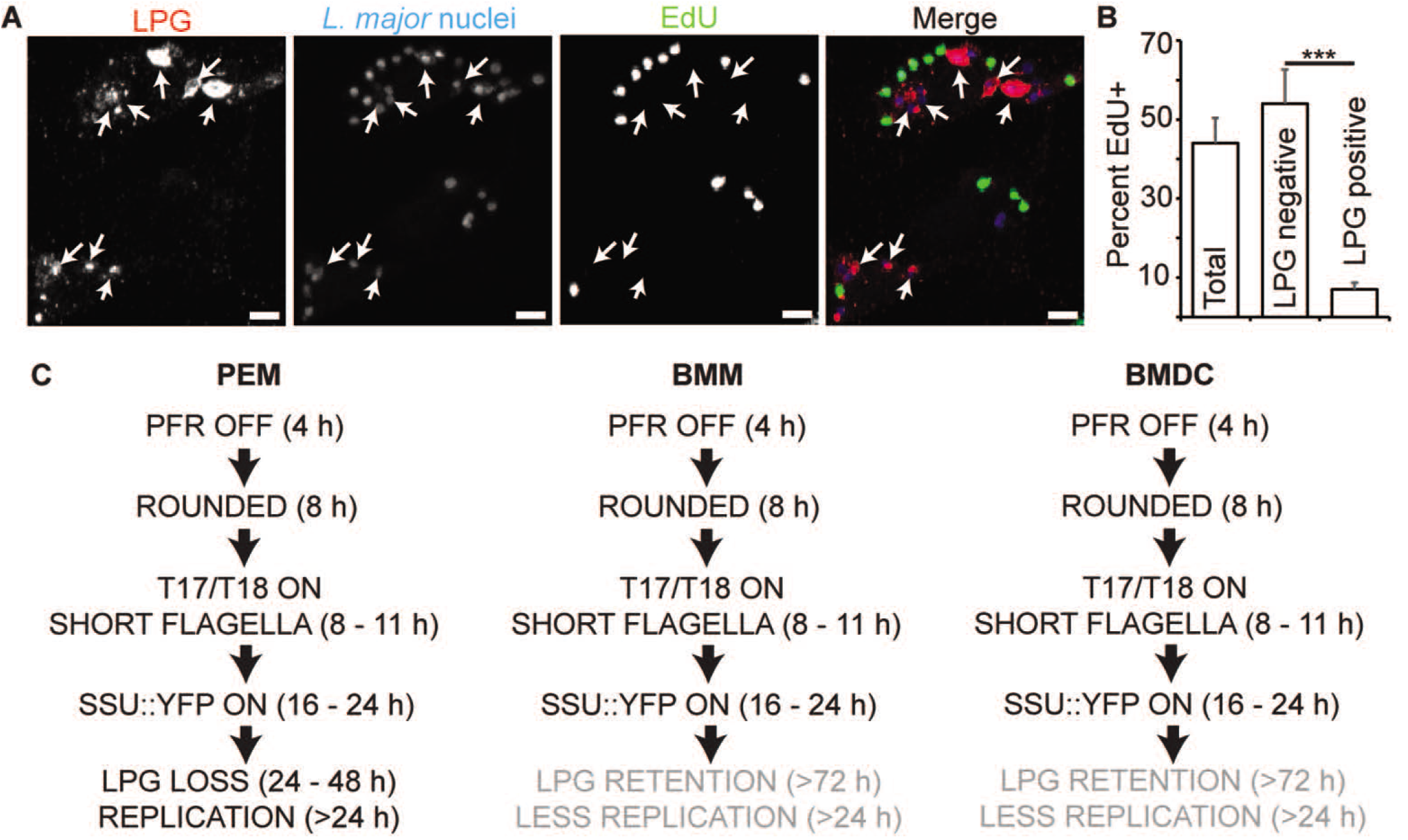
LPG-retaining parasites within DCs 72 hours post-infection have not undergone DNA synthesis. **(A, B)** Parasites were cultured in the presence of thymidine analogue (EdU) for 72 hours following infection of BMDC. Cells were stained to detect EdU-incorporation (green), LPG (red) and parasite nuclei (blue). (A) Representative image of parasites within DCs stained as above. Arrows, parasites showing LPG-positivity. Scale bar, 5 µm. (B) Percent of parasites within DCs 72 hours post-infection that are positive for EdU-incorporation. Data shown include the percent of total parasites that are EdU^+^, as well as the EdU-positivity of parasites that are either LPG-negative or LPG-positive. Data, means ± S.E., n = 3 experiments, ****, *P* < 0.0001 (Chi-square; N = 1185 parasites). **(C)** Comparison of *L. major* amastigogenesis marker transitions, LPG loss, and cell cycle re-entry following infection of PEM, BMM, and BMDC. Overall, the acquisition of early amastigote characteristics is the same in the three host cell types, but LPG loss and DNA replication are reduced or substantially delayed following infection of BMM and BMDC.

### Localization of T17 and T18 epitopes in amastigotes infecting macrophages in vitro

Since the identity of the molecular epitopes recognized by antibodies T17 or T18 were unknown, we attempted to identify them in parasites after induction of *in vitro* amastigogenesis (Fig. S3), via pull-down and mass spectrometry, ultimately without success (data not shown). Amongst many challenges, one possibility is that these antibodies may recognize glycans or other non-protein moieties. We then performed confocal and immuno-electron microscopy to define their localization within amastigotes infecting PEM *in vitro*.

#### T17

By light microscopy, the T17 antibody primarily labelled the amastigote flagellum, with some signal present on the parasite periphery (Fig. 1, 2^nd^ row; Fig. 7A). Examination of T17 immuno-gold labeled *L. major*-infected PEM revealed labelling of the membrane surrounding the amastigote flagellum as well as the flagellum itself (Fig. 7B, C). T17 reactivity also appeared concentrated at the junction between the parasite’s apical end and the phagolysosome membrane (Fig. 7B, D). Remarkably, in these experiments we observed punctate T17 staining within the macrophage cytoplasm in confocal images (Fig. 7E), which corresponded with T17 immuno-EM labeling of the lumen of vesicular structures within the cytoplasm of infected PEMs (Fig. 7F). These data suggest that the T17 epitope may be secreted by amastigote-stage *L. major,* perhaps by trafficking along the flagellum, and suggestive of a role in macrophage infectivity.

**Figure 7.**
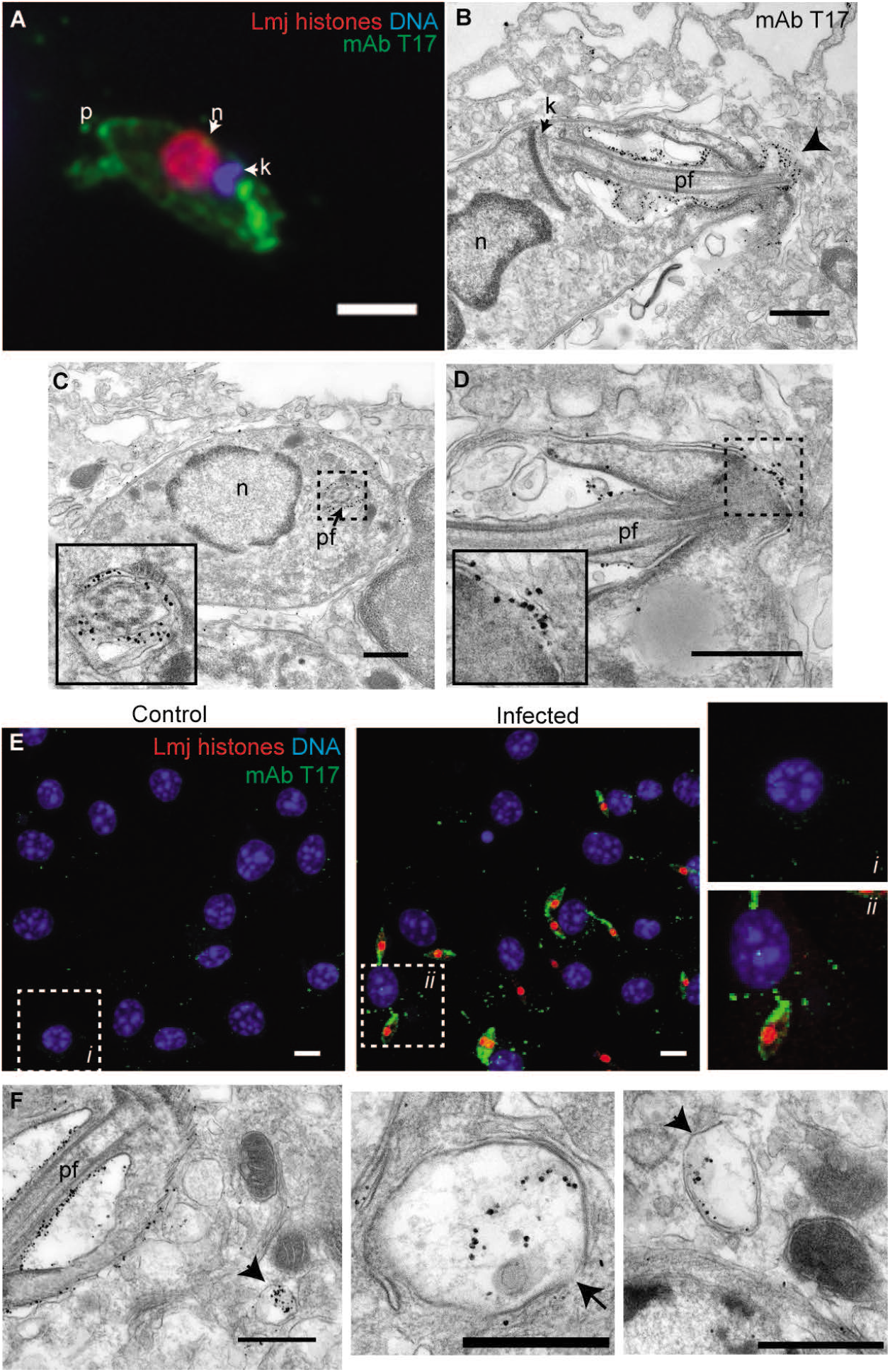
Localization of T17 antibody labeling in amastigotes. **(A)** Epifluorescent microscopy of an amastigote (72 h post metacyclic infection) within a PEM labeled with T17 antibody along with antibodies recognizing parasite histones and DNA stain Hoechst 33342. Scale bar, 2 µm. **(B)** Electron micrograph of an *L. major* amastigote within a PEM following immuno-gold labeling with T17 antibody. Scale bar, 0.5 µm. Arrowhead indicates concentration of T17 immuno-labeling at the distal tip of the amastigote flagellum. **(C)** T17 immuno-labeling around the perimeter of the parasite flagellum. Scale bar, 0.5 µm. **(D)** T17 immuno-labeling is concentrated at the phagolysosome-parasite flagellum contact site, with some signal on the host cell-side of the phagolysosome. Scale bar, 0.5 µm. **(E)** Confocal micrograph of PEM 3 d after *L. major* or mock infection. Samples were stained with T17, antibodies reactive to *Leishmania* histones, and a DNA stain. Zoomed in images of the boxed regions are shown to the right. Scale bar, 10 µm. **(F)** ImmunoEM labeling with T17 of vesicular structures in *L. major*-infected PEM. Images from three different cells shown. Arrows indicate membranous structures containing the parasite antigen recognized by T17 within the PEM cytoplasm. Scale bar, 0.5 µm. Abbreviations: p, parasite posterior; n, nucleus; k, kinetoplast; pf, parasite flagellum.

#### T18

In contrast to antibody T17, by immunofluorescence microscopy antibody T18 recognized an epitope localized to the parasite surface (Fig. 1, row 3), as well as to a region located within the posterior end of the amastigote (furthest from the kinetoplast DNA network; Fig. 8A). These data suggest that T17 and T18 epitopes likely reside on different parasite molecules.

**Figure 8.**
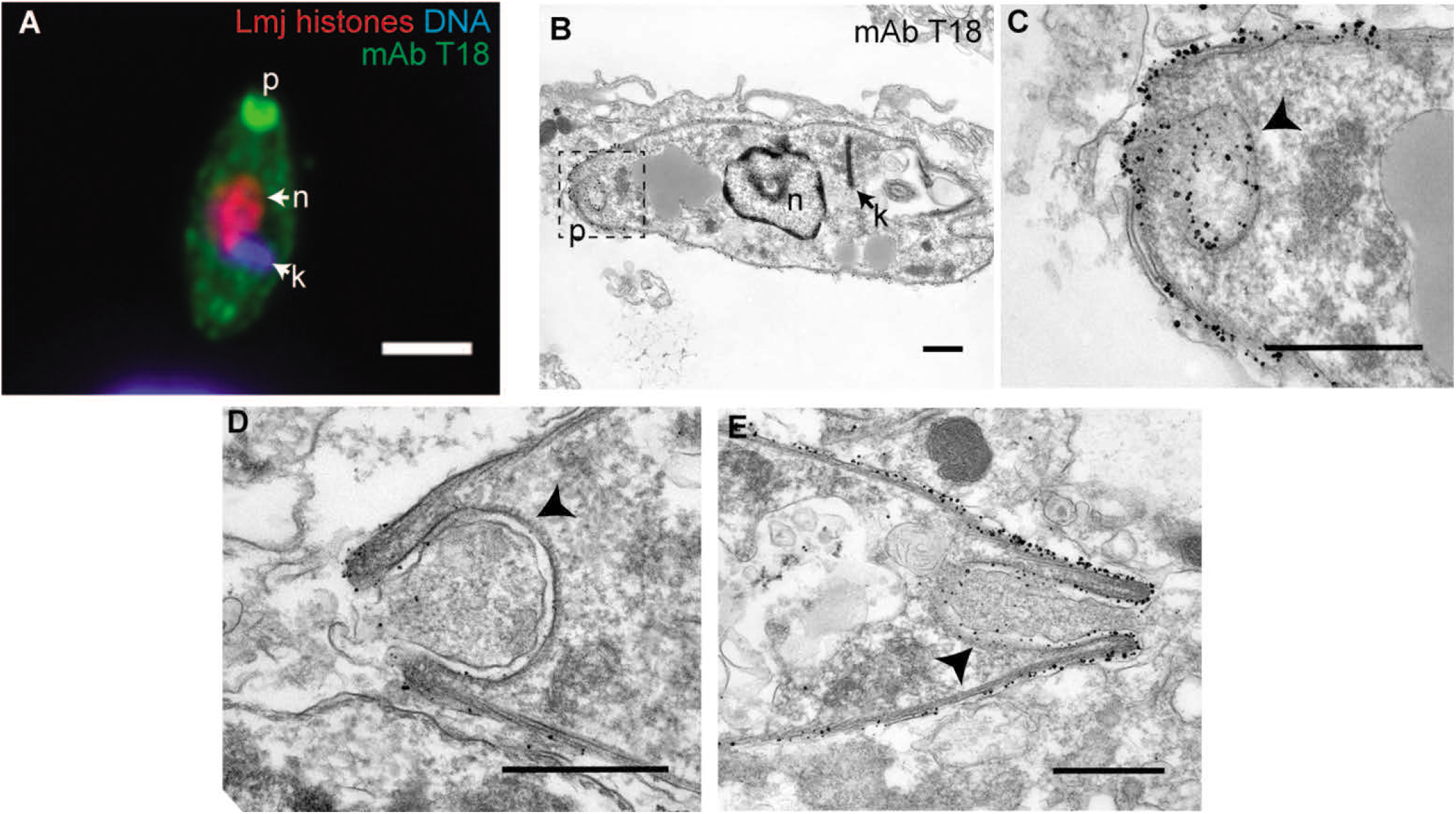
Localization of T18 antibody labeling. **(A)** Epifluorescent microscopy of an amastigote (72 h post metacyclic infection) within a PEM labeled with T18 antibody along with antibodies recognizing parasite histones and DNA stain Hoechst 33342. Scale bar, 2 µm. **(B, C)** Electron micrograph of an *L. major* amastigote within a PEM following immuno-gold labeling with T18 antibody. The region within the dashed box (left) is shown to the right at higher magnification in panel C. **(D, E)** Additional examples of the posterior end of T18 immuno-gold-labeled parasites. **(F)** TEM image of the posterior end of an *L. major* amastigote. These cells were subjected to a different fixation protocol than what was used for immunoEM (see Methods). Arrowheads in panels C-F point to the amastigote-specific structure at the posterior end of the parasite. Scale bar for EM images, 0.5 µm.

By T18 immuno-EM, the posterior tip region showed unexpected morphology, with vesicles and invaginations, with T18 labelling found on the posterior surface and the surface of the invagination (Figs. 8B-E). In many images the invagination lumen appeared to contain material, although whether of parasite or host origin is unknown (Figs. 8C-E). Mitigating concerns about experimental artefacts, the invagination was seen with other fixation techniques as well (Fig. 8F). The nature of the structure(s), and its biological function, if any, are a matter of conjecture. While a similar structure has been previously noted in classic TEM studies of old-world and new-world *Leishmania* amastigotes more than 40 years ago [41–43], it has largely escaped notice and has not previously been defined as being antigenically distinct.

## Discussion

Our study demonstrates that *L. major* parasites can respond to different host cell environments by prolonging their expression of the promastigote stage-specific virulence factor LPG. These studies were enabled by first developing a suite of markers suitable for mapping the process of amastigogenesis at the single cell level, as discussed further below. Interestingly, LPG retention was higher in cell types in which the parasites also showed delayed or reduced initiation of DNA synthesis (BMM and BMDC). Retention of LPG expression is not a result of the parasite’s failure to differentiate into the amastigote stage, as parasites infecting BMM or BMDC clearly did not remain metacyclics and showed nearly identical transitions in terms of several amastigote developmental markers as parasites within highly-permissive PEM.

Our studies identify a population of parasites that resemble amastigotes in terms of lacking PFR expression and being T17, T18, and YFP positive, but that also resemble metacyclics in terms of high-level LPG expression, raising questions as to their nature. Potentially, LPG-loss could represent a “late” marker of amastigogenesis and the PFR^-^ T17^+^ T18^+^ YFP^+^ LPG^+^ cells are an intermediate between the metacyclic and the amastigote stages. However, we identified a number of LPG-positive parasites that had undergone DNA synthesis in BMM and BMDC at 72 hours post infection (Fig. 6B and S5) and in PEM 24 hours post-infection (Fig. S5), indicating that LPG loss is not a prerequisite for cell cycle re-entry. Zilberstein et al have described a ‘maturation’ phase of axenic amastigogenesis that continues to take place even after amastigote-like forms of *L. donovani* have undergone replication [28, 44, 45], and it is possible that the replicative LPG-positive *L. major* are a manifestation of this. Alternatively, their studies differ from those presented here in that they were initiated with replicating log phase promastigotes, rather than purified infective stage metacyclic parasites. A final possibility is that the parasites have completed amastigogenesis yet respond to unfavorable environmental cues by retaining LPG virulence factor expression.

### Significance of LPG persistence

Although none of the host cell types used in this study (PEMs, BMMs, BMDCs) directly mimic any of the various cell types likely to be encountered by metacyclic-stage *Leishmania* following sand fly transmission (e.g. tissue resident dermal macrophages, neutrophils, Langerhans cells, or others), they represent common experimental models which did allow us achieve our goal of exploring the diversity of parasite responses to entry of different host cell types. While amastigote marker transitions were similar regardless of host cell type, we found that parasites display a continuum of responses to different host cell environments in terms of the timing of cell cycle reentry and the expression of the LPG virulence factor, ranging from rapid resumption of replication and LPG loss in some cell types to slower resumption of replication and LPG retention in others. Importantly, these LPG^+^ parasites likely have biologically relevant levels of LPG, as quantitation demonstrated that they express comparable amounts of LPG to metacyclic-stage parasites on a per-cell basis.

Retention of LPG, and potentially other “early” virulence factors that act in parasite establishment, may have important effects on the infected hosts. LPG has roles in parasite survival in macrophages and neutrophils, protection against complement-mediated lysis, oxidant avoidance, cytokine production and the inhibition of phagolysosomal fusion [5, 17, 19–23, 46, 47]. LPG retention would serve to prolong its actions and remodeling the host cell environment, and impacting innate and acquired immune responses susceptible to LPG inhibition [21]. For example, prolonged expression of LPG within DCs as shown here could dramatically impact antigen processing and presentation as well as cytokine production and T-cell activation [46].

Additionally, LPG retention may be beneficial to the parasites as they are transferred from the first host cell infected to the second, through one of several ‘Trojan horse’ routes. In addition to tissue resident macrophages, invading metacyclic-stage parasites are commonly observed infecting neutrophils and dendritic cells *in vivo*, followed by subsequent transfer to macrophages within 24-48 hours after infection [6–8, 48]. In BMM and BMDC, many parasites retained high levels of LPG expression up to 72 hours post infection. As this is well beyond the estimated transit time of *Leishmania* through “first contact” cells to macrophages (∼24 hours), these data establish that the destination macrophages likely encounter a significant number of *Leishmania* retaining biologically relevant levels of LPG capable of mediating virulence functions. For neutrophils, persistence of LPG expression is a necessary step for survival and likely transit [23, 49].

How this transfer of the parasite from the first cell type infected to macrophages occurs is an area of intense study; parasites could be released from the infected cell and subsequently taken up [7] and/or the infected cell itself is engulfed [8, 12, 23, 50], with both routes seeming likely to transfer LPG-retaining parasites. In the future it would be interesting to extend the quantitative single cell approaches developed here to the question of LPG retention in other cell types (neutrophils, inflammatory monocytes, tissue resident macrophages, keratinocytes or other cell types known to harbor *Leishmania* during establishment [6–11]) *in vitro* or *in vivo*. Testing the role of LPG in these widely varying settings will require the application of technologies in which parasite glycoconjugate expression can be manipulated over a short time frame, while the parasite is within the phagolysosome or in transit [51].

### A marker suite for the study of amastigogenesis in *Leishmania major*

Although the primary aim of our study was to determine whether and how *Leishmania* respond to the differing intracellular environments presented by different host cell types, we also developed or characterized a marker suite enabling visualization of the differentiation process of amastigotigenesis at the single cell level that will be generally applicable. Some of the developmental changes associated with amastigogenesis, such as the loss of PFR expression, happened very rapidly upon parasite infection of these cells, with more than 90% of the parasites displaying a PFR^-^ phenotype by 4 hours post-infection. It is well known that the *Leishmania* flagellum undergoes dramatic changes during amastigogenesis [52, 53]. Our results from macrophage-infecting *L. major* are in good agreement with a study tracing flagellar remodeling in *L. mexicana* undergoing amastigotigenesis under host-free axenic conditions *in vitro* [37], with both studies reporting flagellar shortening and PFR loss following host cell infection. Other developmental changes as assessed by amastigote markers occurred later, with T17 and T18 reactivity being the next “markers” to be induced, followed by the induction of YFP fluorescence. Despite the fact that the nature of the antigens recognized by T17 and T18 and the mechanism of YFP down-regulation in metacyclic-stage parasites have not been defined, these still proved to be excellent temporal, cell-based ‘digital’ markers of amastigogenesis. The findings that the induction of amastigote markers appears to occur in an ordered manner (Fig. 2) that is largely independent of host cell type (Fig. 3) and can even occur in the absence of host cell entry (Fig. S3) is overall consistent with the sequential progression of a set of ‘core amastigogenesis’ gene expression and morphological changes seen by microarray studies as *L. donovani* differentiates in axenic culture [25-27, 29, 45, 54]. These changes preceded reentry into the cell cycle, as they had largely gone to completion by 24 hours post infection, a time point at which only 8% of the parasites were BrdU^+^.

### Cellular localization of amastigote-stage markers

Our search for amastigote stage markers led to the selection of the epitopes recognized by monoclonal antibodies T17 and T18 which fulfilled this goal perfectly (Fig. 1). While the identification of the antigens recognized by these antibodies was not required for their use as amastigogenesis markers, it did establish their independence, and their serendipitously interesting cellular localization patterns described below motivated us to undertake further study. Unfortunately, our efforts to identify these antibodies’ molecular targets through combined proteomic and genetic approaches were unsuccessful.

For the epitope recognized by antibody T17, immuno-EM labelling show surface localization to the amastigote flagellum, concentration at the flagellar-parasitophorous vacuole interface, and presence within vesicular structures in the host cell cytoplasm (Fig. 1, Fig. 7). This is interesting when viewed through proposed roles of the amastigote flagellum as a sensory organelle and potentially as a vehicle for delivering virulence factors to or across the phagolysosomal membrane [52]. Indeed, the T17 epitope’s likely path within the infected macrophage would fit this model well.

For the epitope recognized by antibody T18, immunofluorescence and immuno-EM studies showed it to localize to the parasite surface, positioning it to potentially interact with host defenses within the parasitophorous vacuole (Fig. 1). Additionally, the T18 epitope showed strong localization to the parasite’s posterior end, which displayed posterior tip vesicles and invaginations, whose lumen appeared to contain material of undetermined origin (Figs. 8B-E).

The cup-like structure seen at the parasite’s posterior end has previously been noted by electron microscopy of amastigotes from a few *Leishmania* species several decades ago [41–43], but this observation received little attention in the field since no molecular marker for this structure was known prior to our work. However, recent genome wide surveys of protein localization in cultured African trypanosomes have revealed at least 36 candidates with significant posterior location, most of which have *Leishmania* orthologs [55]. Several of these play roles in the parasite cytoskeleton, but many are of unknown function. Interestingly, *L. mexicana* and *L. amazonensis* amastigotes were shown to sequester host proteins including MHC II molecules at a site near their posterior ends [56], possibly connecting the under-studied amastigote posterior structure labeled by T18 with new, interesting biology. Regardless, future studies will be needed to both identify the T18 epitope and dissect its potential roles within the parasite, if any.

In summary, our study traced amastigote development in dendritic cells and two different types of macrophages. We found that overall parasite differentiation was comparable in the three cell types, yet the timing of reentry into the cell cycle and down-regulation of the promastigote-specific virulence factor LPG differed depending on the host cell type (Fig. 6C). This variability in parasite behavior could reflect *Leishmania* responses to the different host cell environments encountered by the parasites early in infection. Finally, the amastigote markers developed in our study will be important tools that can help lay the groundwork for future studies aimed at better understanding the process of amastigogenesis and its regulation.

## Materials and Methods

### Parasite strains and culture

*Leishmania major* Friedlin V1 strain (MHOM/IL/80/Friedlin; abbreviated as LmjF) parasites were grown at 26°C in M199 medium (US Biologicals) supplemented with 40 mM 4-(2-hydroxyethyl)-1-piperazine-ethanesulfonic acid (HEPES) pH 7.4, 50 μM adenosine, 1 μg ml^−1^ biotin, 5 μg ml^−1^ hemin, 2 μg ml^−1^ biopterin and 10% (v/v) heat-inactivated fetal calf serum [51], in some cases containing selective drugs. LmjF parasites expressing YFP (yellow fluorescent protein; *SSU:IR1PHLEO-YFP*) were described previously [51]. *L. major* LV39cl5 Δ*lpg1^-^* [18] was cultured in the above media supplemented with 2 mM L-glutamine, 9 µg ml^-1^ folate and RPMI Vitamin Mix (Sigma). Infective metacyclic-stage parasites were recovered using the density gradient centrifugation method [36]. Prior to infection of host cells, purified metacyclic-stage parasites were opsonized with serum from C5-deficient mice as described [57]. For axenic amastigogenesis experiments shown in Fig. S3, day 3-stationary phase parasites were pelleted and resuspended in warm (37° C) RPMI 1640 media containing L-glutamine and then cultured for 24 h at 37° C in CO_2_ incubator.

### Mammalian host cells

#### Ethics statement

Animal handling and experimental procedures were carried out in strict accordance with the recommendations in the Guide for the Care and Use of Laboratory Animals of the United States National Institutes of Health. Animal studies were approved by the Animal Studies Committee at Washington University (protocol #20-0396) in accordance with the Office of Laboratory Animal Welfare’s guidelines and the Association for Assessment and Accreditation of Laboratory Animal Care International.

Cells were isolated from female C57Bl/6J mice (6-10 weeks old; Jackson Labs). Peritoneal macrophages (PEMs) were elicited by a peritoneal injection of potato starch and harvested and maintained in DMEM (Invitrogen) containing 10% FCS and 2 mM L-glutamine as described [58]. Bone marrow-derived macrophages (BMMs) and dendritic cells (DCs) were harvested as described previously [59]. Briefly, bone marrow was flushed from the femurs of mice and cultured in dendritic cell or macrophage growth media at 37° for 6 days. DCs were cultured in RPMI media without L-glutamine (Gibco) supplemented with 10% fetal calf serum (Hyclone), Glutamax (Gibco), Na pyruvate, non-essential amino acids, and kanamycin (DC media) with the addition of 2% GM-CSF. BMMs were cultured in DMEM (Gibco) supplemented with 10% fetal calf serum, 5% horse serum, Glutamax (Gibco), Na pyruvate, non-essential amino acids, and kanamycin (as described above for the macrophage media) with the addition of 30% L-cell media as the source of M-CSF. For infections, cells were cultured in DC or macrophage media without growth factors. Prior to infection, PEMs, DCs, and BMMs were adhered to sterile glass coverslips in 24 well dishes overnight.

### Infection of host cells

Parasites were added to host cells at a ratio of 5:1. Typically, extracellular parasites were removed by extensive washing 2 hours after parasites were added to host cells. Infected cells were maintained in media which was changed daily. For DNA labeling studies, 0.1 mM 5-bromo-2’-deoxyuridine (BrdU; Sigma) or 0.1 mM 5-ethynyl-2’-deoxyuridine (EdU; Life Technologies) was added for the time indicated in the text.

### Mouse infections and tissue sectioning

The methods used to infect mice and harvest tissues for sectioning have been presented [38]. Briefly, C57Bl/6J mice (6-10 weeks old; Jackson Labs) were injected sub-cutaneously with 10^5^ purified metacyclic parasites in the left hind footpad. 2-3 weeks later, mice were euthanized and the infected footpad was fixed with 4% paraformaldehyde, incubated in sucrose solutions, and embedded in O.C.T. reagent (Ted Pella, Inc.). 10 µm sections of the infected tissue were cut using a cryostat and adhered to microscope slides. Tissue sections were then stained as indicated in the text using the immunofluorescent staining protocol described below.

### Antibodies

*L. major* nuclei were detected with a pool of rabbit antibodies raised against *L. major* histones H_2_A, H_2_A_variant_, H_2_B, H_3_, and H_4_ were pooled at a ratio of 3:2:3:3:1 by titer as determined by Western blot and used at a dilution of 1:750 [38, 60]. BrdU was detected with a rat monoclonal antibody (Abcam) used at 10 μg ml^−1^. For dual-labeling experiments involving YFP, Alexafluor488-conjugated rabbit anti-GFP antisera (Invitrogen) was used at a concentration of 8 μg ml^−1^. The amastigote-specific mouse monoclonal antisera T17 and T18 were a gift from Charles Jaffe (Hebrew University) and were diluted 1:400 [34]. Paraflagellar rod (PFR) was detected with the mouse monoclonal antibody L8C4 (provided by Keith Gull), and was used at a dilution of 1:50 [61]. Lipophoshoglycan (LPG) was detected using two different antisera. For most experiments, Gal-substituted LPG was detected with the mouse monoclonal antibody WIC79.3 [39], which was used at a 1:250 dilution. Where specified, “metacyclic LPG”, in which most of the galactose side chains are capped with D-arabinopyranose, was detected with the mouse monoclonal antibody 3F12 [39] used at a 1:100 dilution. Fluorescent secondary antibodies were: Alexafluor488 goat anti-rabbit, Alexafluor555 goat anti-rabbit, Alexafluor633 goat anti-rabbit, Alexafluor488 goat anti-mouse, Alexafluor594 goat anti-mouse, and Alexafluor488 goat anti-rat (Invitrogen, all used at a concentration of 2 μg ml^−1^).

### Immunofluorescence staining and confocal microscopy

Samples were fixed in 4% (w/v) paraformaldehyde in phosphate-buffered saline (PBS) for 10 minutes. Samples were washed in PBS, and then blocked and permeabilized in PBS containing 5% (v/v) normal goat sera (Vector labs) and 0.1% (v/v) Triton-X-100 for 30 min. The samples were then stained with various combinations of primary antibodies (as described in the text) for 1 h. Unbound antibody was then washed off in PBS and primary antibodies were detected with combinations of fluorescent secondary antibodies (as described in the text) for 40 min, followed by a second wash in PBS. In experiments involving BrdU, fixed samples were washed with distilled water prior to a 40 minute incubation in 2 N HCl. Samples were then extensively washed in PBS prior to blocking and permeabilization as described above. Samples were incubated in anti-BrdU antisera for 2 hours. For experiments involving EdU, EdU was labeled according to the manufacturer’s protocol (Life Technologies) prior to antibody labeling.

Following staining, all samples were mounted in ProLong Gold (Invitrogen). All microscopy was performed on a Zeiss 510 META confocal laser scanning microscope or on an Olympus AX70 microscope equipped with a Retiga 2000 digital camera (QImaging). Cutoffs for saturation and background levels were adjusted with Photoshop software (Adobe). Measurements of parasite and flagellar dimensions were performed using LSM Image Browser Software (Zeiss).

### Immuno-electron microscopy

For immunolocalization by transmission electron microscopy, infected cells were fixed in 4% paraformaldehyde/0.05% glutaraldehyde (Polysciences Inc., Warrington, PA) in 100mM PIPES/0.5mM MgCl2, pH 7.2 for 1 hr at 4°C. For ultrastructural analysis, samples were fixed in 2% paraformaldehyde/2.5% glutaraldehyde (Polysciences Inc., Warrington, PA) in 100mM PIPES/0.5mM MgCl2, pH 7.2 for 1 hr at 20°C. After either fixation protocol, samples were then infiltrated overnight in the cryoprotectant 2.3M sucrose/20% polyvinyl pyrrolidone in PIPES/MgCl2 at 4°C. To permeabilize cells for antibody labeling samples were plunge-frozen in liquid nitrogen and subsequently thawed in PBS at room temperature. This technique was confirmed to permeabilize the host cell membrane and intracellular organelle membranes. Samples were probed with the primary antibodies at 1:250 dilutions followed by FluoroNanogold anti-mouse Fab (1:250; Nanoprobes, Yaphank, NY) and silver enhancement (Nanoprobes HQ silver enhancement kit). Samples were washed in phosphate buffer and post-fixed in 1% osmium tetroxide (Polysciences Inc., Warrington, PA) for 1 hr. Samples were then rinsed extensively in dH20 prior to *en bloc* staining with 1% aqueous uranyl acetate (Ted Pella Inc., Redding, CA) for 1 hr. Following several rinses in dH20, samples were dehydrated in a graded series of ethanol and embedded in Eponate 12 resin (Ted Pella Inc.). Sections of 95 nm were cut with a Leica Ultracut UCT ultramicrotome (Leica Microsystems Inc., Bannockburn, IL), stained with uranyl acetate and lead citrate, and viewed on a JEOL 1200 EX transmission electron microscope (JEOL USA Inc., Peabody, MA). All pre-labeling experiments were conducted in parallel with omission of the primary antibody. These controls were consistently negative at the concentration of Nanoprobes-conjugated secondary antibodies used in these studies.

### Quantitation of LPG abundance

Samples were stained to detect parasite histones and PGs and confocal microscopy performed as described above. 3-dimensional confocal image stacks were then compressed into a single 2-dimensional image which was then used for subsequent analysis. For samples harvested less than 24 hours post infection, at which time the parasite’s outline could be visualized with WIC79.3 staining, Volocity software (Improvision) was used to trace the outline of the parasites and then measure the sum of the WIC79.3 (red) intensity within the traced area. For samples harvested 24 hours or later after infection, this method was unusable because the outline of the parasites was increasingly invisible. Thus, we measured the sum of WIC79.3 intensities within a 44.8 µm^2^ circle centered at the parasite nucleus. We did not use this method with parasites at time points prior to 24 hours because a circle is a poor approximation of the elongated shape of metacyclic-stage parasites.

### Flow cytometry

YFP fluorescence was quantitated on a FACSCalibur flow cytometer (BD Biosciences) using the FL1 channel. FSC and SSC were used to gate on the single cell population.

### Data analysis and Statistics

Unless stated otherwise, the data reported throughout the paper is the mean of at least three independent experiments in which >300 parasites were scored per experiment. *P* values < 0.05 were considered significant using either ANOVA followed by a post hoc Bonferroni multiple comparisons test or a Chi-squared test as specified in the text or figure legends.

## Abbreviations

DC: dendritic cell
BrdU: 5-bromo-2-deoxyuridine
LPG: lipophosphoglycan
BMM: bone marrow derived macrophages
PEM: peritoneal macrophages
BMDC: bone-marrow derived dendritic cells
PPG: proteophosphoglycan

## Acknowledgements

We thank Dr. Charles Jaffe for providing the amastigote specific antibodies T17 and T18, and Dr. Keith Gull for providing the anti-PFR2 antibody. This work was funded by NIH R01 AI31078 to S.M.B. We thank Drs. Marco Colonna and Stephen McCartney for assistance with bone marrow-derived cells, Dr. Nathan Peters for discussions concerning LPG persistence and neutrophils, Dr. Igor Almeida for discussions and assistance with immunoprecipitation and mass spectrometry, and Dr. Ziyin Li for discussions concerning ‘posterior’ proteins in African trypanosomes.

## Author contributions

M.A.M and W.L.B performed experiments; M.A.M and S.M.B designed experiments, analyzed results, and wrote and revised the manuscript.

## SUPPLEMENTARY FIGURE LEGENDS

**Figure S1.**
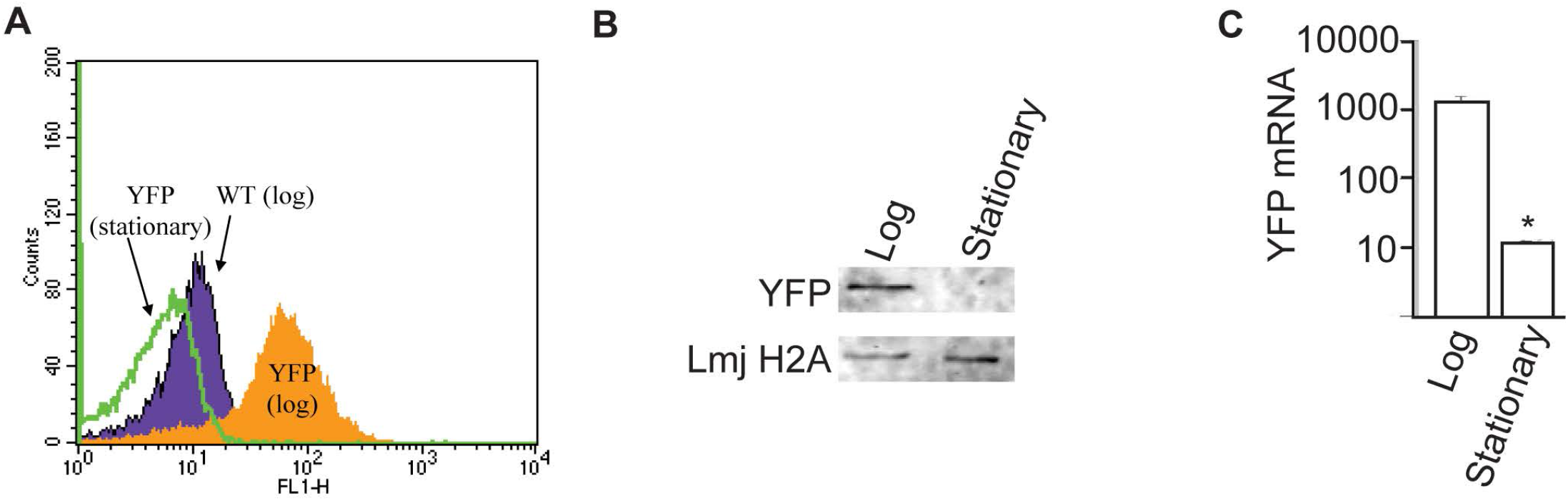
Developmental regulation of YFP reporter. **(A)** Flow cytometric analysis of YFP signal from WT *L. major* and *L. major* expressing YFP from the small ribosomal subunit locus under log-phase (procyclic promastigotes) or stationary-phase (includes metacyclic promastigotes) conditions. **(B)** Western blot analysis of YFP expression in log phase and stationary phase parasites stably transfected with YFP transgene. *L. major* histone H2A levels were assessed as a protein loading control. **(C)** The abundance of YFP mRNA in log and stationary phase parasites was determined by q-RT-PCR. *, *P* < 0.05. Abbreviations: p, parasite posterior; n, nucleus; k, kinetoplast.

**Figure S2.**
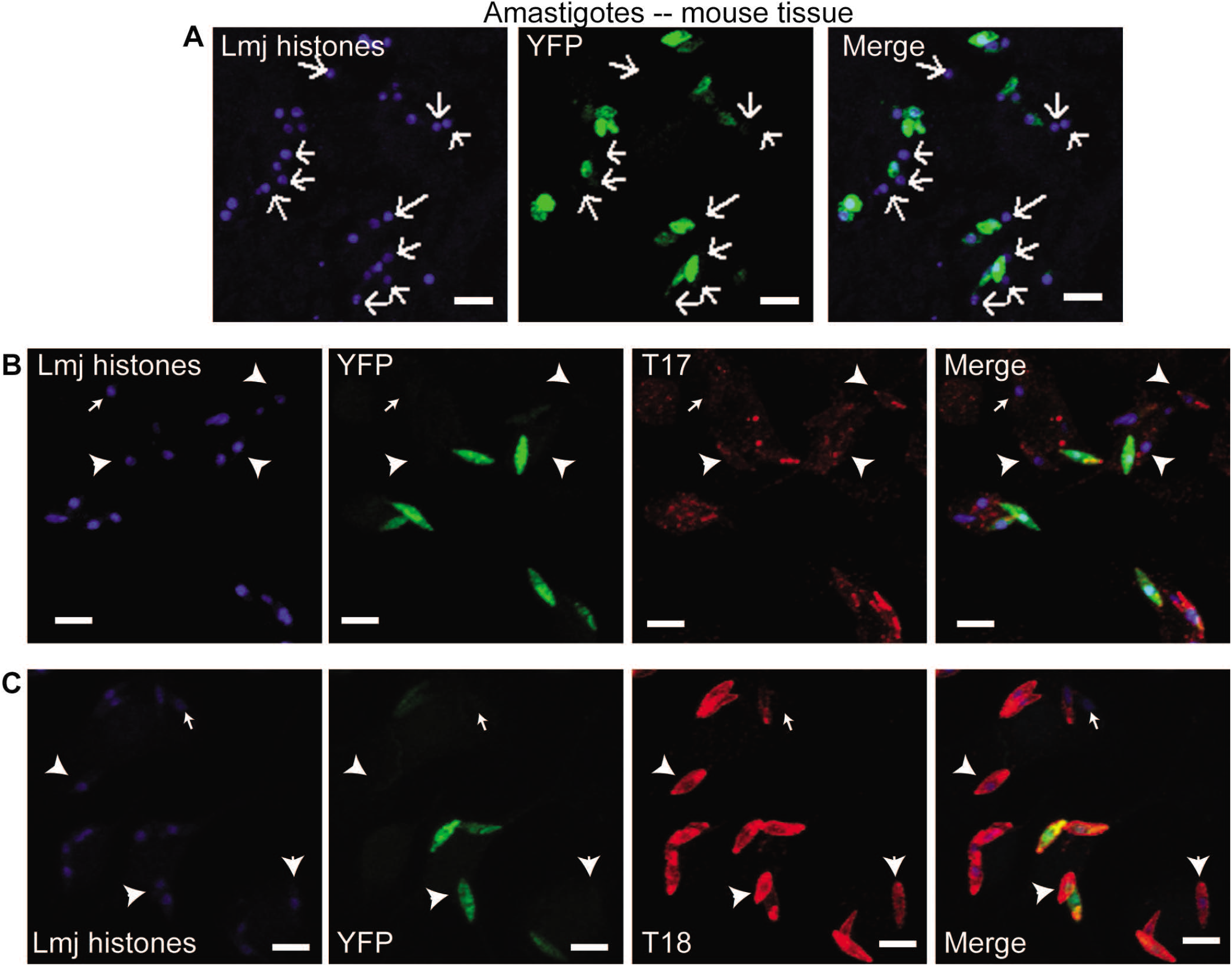
mAb T17 and T18 reactivity precedes YFP expression during metacyclic-to-amastigote transition. **(A)** C57B6 mice were injected subcutaneously in footpad tissue with metacyclic stage parasites stably transfected with YFP transgene. 16 d later, mice were sacrificed and infected footpad tissue sectioned and stained to detect parasite histones (blue) and YFP. YFP-negative parasites are indicated by arrows. Scale bars, 5 µm. **(B, C)** PEMs were fixed 10 h after infection and stained with T17 (B) or T18 (C) antibodies (red). Parasites were identified based on nuclear staining with antibodies against parasite histones (blue). The percent of parasites positive for T17 or T18 reactivity and YFP fluorescence (green) was determined by analysis of confocal micrographs. Arrowheads indicate T17/T18 positive, YFP-negative parasites, and arrows indicate double-negative parasites. Scale bars, 5 µm.

**Figure S3.**
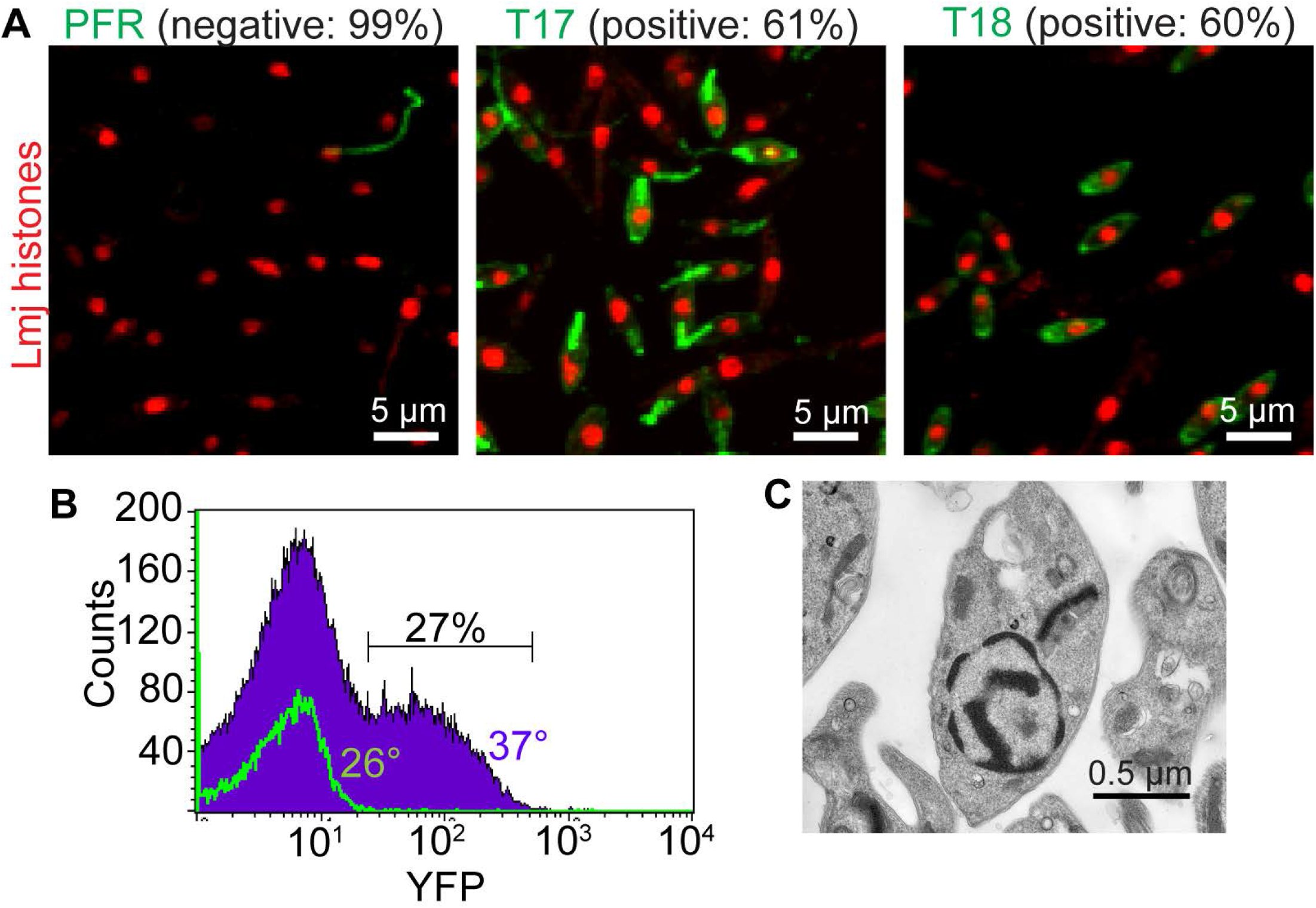
Acquisition of amastigote traits by *L. major* cultured at 37⁰ C in the absence of host cells. *L. major* promastigotes were cultured until the third day of stationary phase at 26⁰ C, then placed in fresh media and cultured at 37⁰ C for 24 hours prior to harvest and analysis. **(A)** Parasites were fixed and stained with the indicated markers prior to confocal microscopy. Micrographs were visually scored for PFR negativity or T17/T18 positivity. N > 200 parasites. **(B)** Flow cytometric analysis of YFP fluorescence of the starting culture of stationary phase parasites (green, 26⁰ C) and parasites following 24 hours of culture at 37⁰ C (purple). **(C)** Representative electron micrograph of a parasite following culture at 37⁰ C that has typical amastigote morphology including a round shape and a spacious flagellar pocket.

**Figure S4.**
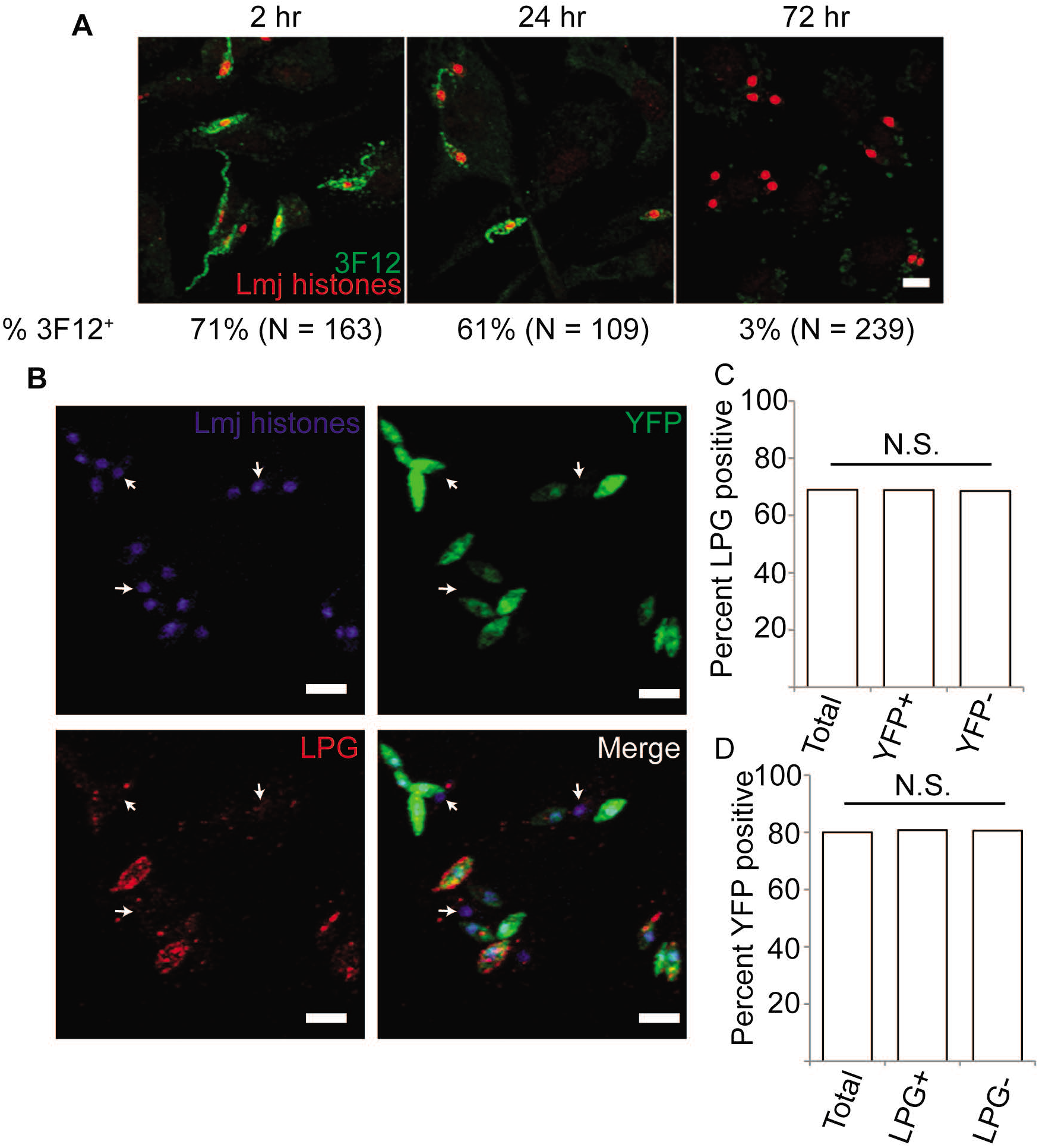
LPG down-regulation is coincident with loss of metacyclic-specific arabinose-capped LPG but independent of YFP induction. **(A)** PEMs were infected with *L. major* and harvested 2, 24, and 72 hours after infection. Ara-capped LPG is detected with mAB 3F12 (green), and parasite histones are shown in red. Data indicate the percent of parasites positive for 3F12 labeling out of the total number of parasites analyzed (N). Scale bar, 5 µm. **(B)** Representative image of parasites within BMM fixed 72 h post infection showing LPG and/or YFP positivity. Parasite nuclei are shown in blue. Arrows indicate LPG and YFP double-negative parasites. Scale bar, 5 µm. Analysis of images like that in (B) was performed to determine the LPG-positivity **(C)** or YFP-positivity **(D)** of total parasites or within the indicated parasite sup-populations. N = 362 parasites. N.S., not specific (Chi-square).

**Figure S5.**
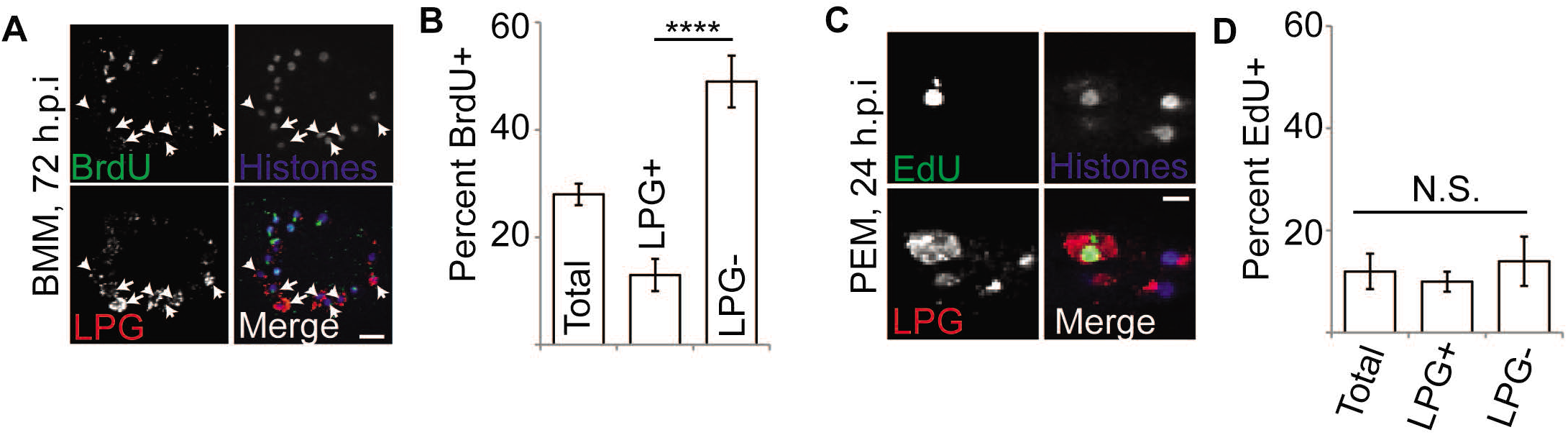
LPG loss is not prerequisite for DNA synthesis. **(A, B)** *L. major*-infected BMM were cultured in the presence of BrdU for 72 hours prior to fixation, immunolabeling, and confocal microscopy. A representative image is shown in (A). Arrows indicate LPG-retaining parasites, which tend to be BrdU-negative as quantitated by Chi-square analysis in (B). Data shown include the percent of total parasites that are BrdU^+^, as well as the BrdU-positivity of parasites that are either LPG-negative or LPG-positive. Data, means ± S.E., n = 3 experiments, ***, *P* < 0.0001 (Chi-square; N = 848 parasites). **(C, D)** PEMs were infected with *L. major* metacyclic stage parasites for 24 h in the presence of EdU prior to fixation and staining to detect EdU (green), LPG (red) and parasite nuclei (blue). A Representative confocal image of a parasite showing double-positive labeling for EdU and LPG along with several EdU-negative parasites is shown in (C) with quantitation of EdU positivity amongst total parasites, LPG-positive, and LPG-negative parasites shown in (D). N.S., not significant per Chi-square analysis as in Fig. 6. N = 429 parasites.

## Notes

### Competing Interest Statement

The authors have declared no competing interest.

### Summary of Updates

this is a revised manuscript

## References

1. Desjeux P. Leishmaniasis. Nature reviews Microbiology. 2004;2(9):692. PubMed PMID: 15378809.

2. Mosser DM, Brittingham A. Leishmania, macrophages and complement: a tale of subversion and exploitation. Parasitology. 1997;115 Suppl:S9–23. PubMed PMID: 9571687.

3. Guimaraes-Costa AB, Nascimento MT, Froment GS, Soares RP, Morgado FN, Conceicao-Silva F, et al. Leishmania amazonensis promastigotes induce and are killed by neutrophil extracellular traps. Proceedings of the National Academy of Sciences of the United States of America. 2009;106(16):6748–53. doi: 10.1073/pnas.0900226106. PubMed PMID: 19346483; PubMed Central PMCID: PMC2672475.

4. Gabriel C, McMaster WR, Girard D, Descoteaux A. Leishmania donovani promastigotes evade the antimicrobial activity of neutrophil extracellular traps. J Immunol. 2010;185(7):4319–27. PubMed PMID: 20826753.

5. Moradin N, Descoteaux A. Leishmania promastigotes: building a safe niche within macrophages. Frontiers in cellular and infection microbiology. 2012;2:121. PubMed PMID: 23050244.

6. Ng LG, Hsu A, Mandell MA, Roediger B, Hoeller C, Mrass P, et al. Migratory dermal dendritic cells act as rapid sensors of protozoan parasites. PLoS Pathog. 2008;4(11):e1000222. PubMed PMID: 19043558.

7. Peters NC, Egen JG, Secundino N, Debrabant A, Kimblin N, Kamhawi S, et al. In vivo imaging reveals an essential role for neutrophils in leishmaniasis transmitted by sand flies. Science. 2008;321(5891):970–4. PubMed PMID: 18703742.

8. Chaves MM, Lee SH, Kamenyeva O, Ghosh K, Peters NC, Sacks D. The role of dermis resident macrophages and their interaction with neutrophils in the early establishment of Leishmania major infection transmitted by sand fly bite. PLoS Pathog. 2020;16(11):e1008674. Epub 2020/11/03. doi: 10.1371/journal.ppat.1008674. PubMed PMID: 33137149; PubMed Central PMCID: PMCPMC7660907.

9. ElHassan AM, Gaafar A, Theander TG. Antigen-presenting cells in human cutaneous leishmaniasis due to Leishmania major. Clinical and experimental immunology. 1995;99(3):445–53. PubMed PMID: 7882568; PubMed Central PMCID: PMC1534205.

10. Bogdan C, Donhauser N, Doring R, Rollinghoff M, Diefenbach A, Rittig MG. Fibroblasts as host cells in latent leishmaniosis. J Exp Med. 2000;191(12):2121–30. PubMed PMID: 10859337.

11. Ribeiro-Gomes FL, Peters NC, Debrabant A, Sacks DL. Efficient capture of infected neutrophils by dendritic cells in the skin inhibits the early anti-leishmania response. PLoS Pathog. 2012;8(2):e1002536. Epub 2012/02/24. doi: 10.1371/journal.ppat.1002536. PubMed PMID: 22359507; PubMed Central PMCID: PMCPMC3280984.

12. Kleinholz CL, Riek-Burchardt M, Seiss EA, Amore J, Gintschel P, Philipsen L, et al. Ly6G deficiency alters the dynamics of neutrophil recruitment and pathogen capture during Leishmania major skin infection. Sci Rep. 2021;11(1):15071. Epub 2021/07/25. doi: 10.1038/s41598-021-94425-9. PubMed PMID: 34302006; PubMed Central PMCID: PMCPMC8302578.

13. Romano A, Carneiro MBH, Doria NA, Roma EH, Ribeiro-Gomes FL, Inbar E, et al. Divergent roles for Ly6C+CCR2+CX3CR1+ inflammatory monocytes during primary or secondary infection of the skin with the intra-phagosomal pathogen Leishmania major. PLoS Pathog. 2017;13(6):e1006479. Epub 2017/07/01. doi: 10.1371/journal.ppat.1006479. PubMed PMID: 28666021; PubMed Central PMCID: PMCPMC5509374.

14. Franco LH, Beverley SM, Zamboni DS. Innate immune activation and subversion of Mammalian functions by leishmania lipophosphoglycan. J Parasitol Res. 2012;2012:165126. doi: 10.1155/2012/165126. PubMed PMID: 22523640; PubMed Central PMCID: PMC3317186.

15. Olivier M, Atayde VD, Isnard A, Hassani K, Shio MT. Leishmania virulence factors: focus on the metalloprotease GP63. Microbes Infect. 2012;14(15):1377–89. doi: 10.1016/j.micinf.2012.05.014. PubMed PMID: 22683718.

16. Beverley SM, Turco SJ. Lipophosphoglycan (LPG) and the identification of virulence genes in the protozoan parasite Leishmania. Trends Microbiol. 1998;6(1):35–40. PubMed PMID: 9481823.

17. Spath GF, Garraway LA, Turco SJ, Beverley SM. The role(s) of lipophosphoglycan (LPG) in the establishment of Leishmania major infections in mammalian hosts. Proceedings of the National Academy of Sciences of the United States of America. 2003;100(16):9536–41. PubMed PMID: 12869694.

18. Spath GF, Epstein L, Leader B, Singer SM, Avila HA, Turco SJ, et al. Lipophosphoglycan is a virulence factor distinct from related glycoconjugates in the protozoan parasite Leishmania major. Proceedings of the National Academy of Sciences of the United States of America. 2000;97(16):9258–63. PubMed PMID: 10908670.

19. Lodge R, Descoteaux A. Modulation of phagolysosome biogenesis by the lipophosphoglycan of Leishmania. Clin Immunol. 2005;114(3):256–65. Epub 2005/02/22. doi: 10.1016/j.clim.2004.07.018. PubMed PMID: 15721836.

20. Olivier M, Gregory DJ, Forget G. Subversion mechanisms by which Leishmania parasites can escape the host immune response: a signaling point of view. Clin Microbiol Rev. 2005;18(2):293–305. PubMed PMID: 15831826.

21. Liu D, Kebaier C, Pakpour N, Capul AA, Beverley SM, Scott P, et al. Leishmania major phosphoglycans influence the host early immune response by modulating dendritic cell functions. Infect Immun. 2009;77(8):3272–83. PubMed PMID: 19487470.

22. Lodge R, Diallo TO, Descoteaux A. Leishmania donovani lipophosphoglycan blocks NADPH oxidase assembly at the phagosome membrane. Cellular microbiology. 2006;8(12):1922–31. PubMed PMID: 16848789.

23. Gueirard P, Laplante A, Rondeau C, Milon G, Desjardins M. Trafficking of Leishmania donovani promastigotes in non-lytic compartments in neutrophils enables the subsequent transfer of parasites to macrophages. Cellular microbiology. 2008;10(1):100–11. PubMed PMID: 17651446.

24. de Carvalho RVH, Andrade WA, Lima-Junior DS, Dilucca M, de Oliveira CV, Wang K, et al. Leishmania Lipophosphoglycan Triggers Caspase-11 and the Non-canonical Activation of the NLRP3 Inflammasome. Cell reports. 2019;26(2):429–37 e5. Epub 2019/01/10. doi: 10.1016/j.celrep.2018.12.047. PubMed PMID: 30625325; PubMed Central PMCID: PMCPMC8022207.

25. Leifso K, Cohen-Freue G, Dogra N, Murray A, McMaster WR. Genomic and proteomic expression analysis of Leishmania promastigote and amastigote life stages: the Leishmania genome is constitutively expressed. Molecular and biochemical parasitology. 2007;152(1):35–46. PubMed PMID: 17188763.

26. McNicoll F, Drummelsmith J, Muller M, Madore E, Boilard N, Ouellette M, et al. A combined proteomic and transcriptomic approach to the study of stage differentiation in Leishmania infantum. Proteomics. 2006;6(12):3567–81. PubMed PMID: 16705753.

27. Akopyants NS, Matlib RS, Bukanova EN, Smeds MR, Brownstein BH, Stormo GD, et al. Expression profiling using random genomic DNA microarrays identifies differentially expressed genes associated with three major developmental stages of the protozoan parasite Leishmania major. Molecular and biochemical parasitology. 2004;136(1):71–86. PubMed PMID: 15138069.

28. Saxena A, Lahav T, Holland N, Aggarwal G, Anupama A, Huang Y, et al. Analysis of the Leishmania donovani transcriptome reveals an ordered progression of transient and permanent changes in gene expression during differentiation. Molecular and biochemical parasitology. 2007;152(1):53–65. PubMed PMID: 17204342.

29. Srividya G, Duncan R, Sharma P, Raju BV, Nakhasi HL, Salotra P. Transcriptome analysis during the process of in vitro differentiation of Leishmania donovani using genomic microarrays. Parasitology. 2007;134(Pt 11):1527–39. PubMed PMID: 17553180.

30. Alcolea PJ, Alonso A, Gomez MJ, Postigo M, Molina R, Jimenez M, et al. Stage-specific differential gene expression in Leishmania infantum: from the foregut of Phlebotomus perniciosus to the human phagocyte. BMC genomics. 2014;15:849. doi: 10.1186/1471-2164-15-849. PubMed PMID: 25281593; PubMed Central PMCID: PMC4203910.

31. Rochette A, Raymond F, Ubeda JM, Smith M, Messier N, Boisvert S, et al. Genome-wide gene expression profiling analysis of Leishmania major and Leishmania infantum developmental stages reveals substantial differences between the two species. BMC genomics. 2008;9:255. doi: 10.1186/1471-2164-9-255. PubMed PMID: 18510761; PubMed Central PMCID: PMC2453527.

32. Moore LL, Santrich C, LeBowitz JH. Stage-specific expression of the Leishmania mexicana paraflagellar rod protein PFR-2. Molecular and biochemical parasitology. 1996;80(2):125–35. PubMed PMID: 8892290.

33. Mishra KK, Holzer TR, Moore LL, LeBowitz JH. A negative regulatory element controls mRNA abundance of the Leishmania mexicana Paraflagellar rod gene PFR2. Eukaryotic cell. 2003;2(5):1009–17. PubMed PMID: 14555483; PubMed Central PMCID: PMC219351.

34. Jaffe CL, Rachamim N. Amastigote stage-specific monoclonal antibodies against Leishmania major. Infect Immun. 1989;57(12):3770–7. PubMed PMID: 2680982.

35. Zakai HA, Chance ML, Bates PA. In vitro stimulation of metacyclogenesis in Leishmania braziliensis, L. donovani, L. major and L. mexicana. Parasitology. 1998;116 (Pt 4):305–9. PubMed PMID: 9585932.

36. Spath GF, Beverley SM. A lipophosphoglycan-independent method for isolation of infective Leishmania metacyclic promastigotes by density gradient centrifugation. Experimental parasitology. 2001;99(2):97–103. PubMed PMID: 11748963.

37. Wheeler RJ, Gluenz E, Gull K. Basal body multipotency and axonemal remodelling are two pathways to a 9+0 flagellum. Nature communications. 2015;6:8964. doi: 10.1038/ncomms9964. PubMed PMID: 26667778; PubMed Central PMCID: PMC4682162.

38. Mandell MA, Beverley SM. Continual renewal and replication of persistent Leishmania major parasites in concomitantly immune hosts. Proceedings of the National Academy of Sciences of the United States of America. 2017;114(5):E801–E10. doi: 10.1073/pnas.1619265114. PubMed PMID: 28096392; PubMed Central PMCID: PMC5293024.

39. Kelleher M, Bacic A, Handman E. Identification of a macrophage-binding determinant on lipophosphoglycan from Leishmania major promastigotes. Proceedings of the National Academy of Sciences of the United States of America. 1992;89(1):6–10. PubMed PMID: 1370357.

40. Salic A, Mitchison TJ. A chemical method for fast and sensitive detection of DNA synthesis in vivo. Proceedings of the National Academy of Sciences of the United States of America. 2008;105(7):2415–20. PubMed PMID: 18272492.

41. Gardener PJ. Pellicle-associated structures in the amastigote stage of Trypanosoma cruzi and Leishmania species. Ann Trop Med Parasitol. 1974;68(2):167–76. Epub 1974/06/01. doi: 10.1080/00034983.1974.11686935. PubMed PMID: 4212227.

42. Pan AA, Pan SC. Leishmania mexicana: comparative fine structure of amastigotes and promastigotes in vitro and in vivo. Experimental parasitology. 1986;62(2):254–65. Epub 1986/10/01. doi: 10.1016/0014-4894(86)90030-5. PubMed PMID: 3743717.

43. Pham TD, Azar HA, Moscovic EA, Kurban AK. The ultrastructure of Leishmania tropica in the oriental sore. Ann Trop Med Parasitol. 1970;64(1):1–4. Epub 1970/03/01. doi: 10.1080/00034983.1970.11686657. PubMed PMID: 5485707.

44. Tsigankov P, Gherardini PF, Helmer-Citterich M, Zilberstein D. What has proteomics taught us about Leishmania development? Parasitology. 2012;139(9):1146–57. Epub 2012/03/01. doi: 10.1017/S0031182012000157. PubMed PMID: 22369930.

45. Barak E, Amin-Spector S, Gerliak E, Goyard S, Holland N, Zilberstein D. Differentiation of Leishmania donovani in host-free system: analysis of signal perception and response. Molecular and biochemical parasitology. 2005;141(1):99–108. Epub 2005/04/07. doi: 10.1016/j.molbiopara.2005.02.004. PubMed PMID: 15811531.

46. Liu D, Okwor I, Mou Z, Beverley SM, Uzonna JE. Deficiency of Leishmania phosphoglycans influences the magnitude but does not affect the quality of secondary (memory) anti-Leishmania immunity. PLoS One. 2013;8(6):e66058. Epub 2013/06/19. doi: 10.1371/journal.pone.0066058. PubMed PMID: 23776605; PubMed Central PMCID: PMCPMC3679009.

47. Favila MA, Geraci NS, Jayakumar A, Hickerson S, Mostrom J, Turco SJ, et al. Differential Impact of LPG-and PG-Deficient Leishmania major Mutants on the Immune Response of Human Dendritic Cells. PLoS neglected tropical diseases. 2015;9(12):e0004238. Epub 2015/12/03. doi: 10.1371/journal.pntd.0004238. PubMed PMID: 26630499; PubMed Central PMCID: PMCPMC4667916.

48. Thalhofer CJ, Chen Y, Sudan B, Love-Homan L, Wilson ME. Leukocytes infiltrate the skin and draining lymph nodes in response to the protozoan Leishmania infantum chagasi. Infect Immun. 2011;79(1):108–17. PubMed PMID: 20937764.

49. Quintela-Carvalho G, Goicochea AMC, Mancur-Santos V, Viana SM, Luz YDS, Dias BRS, et al. Leishmania infantum Defective in Lipophosphoglycan Biosynthesis Interferes With Activation of Human Neutrophils. Frontiers in cellular and infection microbiology. 2022;12:788196. Epub 2022/04/26. doi: 10.3389/fcimb.2022.788196. PubMed PMID: 35463648; PubMed Central PMCID: PMCPMC9019130.

50. Ritter U, Frischknecht F, van Zandbergen G. Are neutrophils important host cells for Leishmania parasites? Trends Parasitol. 2009;25(11):505–10. PubMed PMID: 19762280.

51. Madeira da Silva L, Owens KL, Murta SM, Beverley SM. Regulated expression of the Leishmania major surface virulence factor lipophosphoglycan using conditionally destabilized fusion proteins. Proceedings of the National Academy of Sciences of the United States of America. 2009;106(18):7583–8. PubMed PMID: 19383793.

52. Gluenz E, Hoog JL, Smith AE, Dawe HR, Shaw MK, Gull K. Beyond 9+0: noncanonical axoneme structures characterize sensory cilia from protists to humans. Faseb J. 2010;24(9):3117–21. PubMed PMID: 20371625.

53. Landfear SM. New Vistas in the Biology of the Flagellum-Leishmania Parasites. Pathogens. 2022;11(4). Epub 2022/04/24. doi: 10.3390/pathogens11040447. PubMed PMID: 35456123; PubMed Central PMCID: PMC9024700.

54. Holzer TR, McMaster WR, Forney JD. Expression profiling by whole-genome interspecies microarray hybridization reveals differential gene expression in procyclic promastigotes, lesion-derived amastigotes, and axenic amastigotes in Leishmania mexicana. Molecular and biochemical parasitology. 2006;146(2):198–218. PubMed PMID: 16430978.

55. Billington K, Halliday C, Madden R, Dyer P, Carrington M, Vaughan S, et al. Genome-wide subcellular protein localisation in the flagellate parasite *Trypanosoma brucei*. bioRxiv. 2022:2022.06.09.495287. doi: 10.1101/2022.06.09.495287.

56. Antoine JC, Lang T, Prina E, Courret N, Hellio R. H-2M molecules, like MHC class II molecules, are targeted to parasitophorous vacuoles of Leishmania-infected macrophages and internalized by amastigotes of L. amazonensis and L. mexicana. Journal of cell science. 1999;112 (Pt 15):2559–70. Epub 1999/07/08. doi: 10.1242/jcs.112.15.2559. PubMed PMID: 10393812.

57. Spath GF, Lye LF, Segawa H, Sacks DL, Turco SJ, Beverley SM. Persistence without pathology in phosphoglycan-deficient Leishmania major. Science. 2003;301(5637):1241–3. PubMed PMID: 12947201.

58. Capul AA, Hickerson S, Barron T, Turco SJ, Beverley SM. Comparisons of mutants lacking the Golgi UDP-galactose or GDP-mannose transporters establish that phosphoglycans are important for promastigote but not amastigote virulence in Leishmania major. Infect Immun. 2007;75(9):4629–37. PubMed PMID: 17606605.

59. Gitlin L, Barchet W, Gilfillan S, Cella M, Beutler B, Flavell RA, et al. Essential role of mda-5 in type I IFN responses to polyriboinosinic:polyribocytidylic acid and encephalomyocarditis picornavirus. Proceedings of the National Academy of Sciences of the United States of America. 2006;103(22):8459–64. PubMed PMID: 16714379.

60. Anderson BA, Wong IL, Baugh L, Ramasamy G, Myler PJ, Beverley SM. Kinetoplastid-specific histone variant functions are conserved in Leishmania major. Molecular and biochemical parasitology. 2013;191(2):53–7. Epub 2013/10/02. doi: S0166-6851(13)00136-9 [pii] 10.1016/j.molbiopara.2013.09.005. PubMed PMID: 24080031; PubMed Central PMCID: PMC3863619.

61. Kohl L, Sherwin T, Gull K. Assembly of the paraflagellar rod and the flagellum attachment zone complex during the Trypanosoma brucei cell cycle. The Journal of eukaryotic microbiology. 1999;46(2):105–9. PubMed PMID: 10361731.

